# One Cell At a Time: A Unified Framework to Integrate and Analyze Single-cell RNA-seq Data

**DOI:** 10.1101/2021.05.12.443814

**Authors:** Chloe X. Wang, Lin Zhang, Bo Wang

## Abstract

The surge of single-cell RNA sequencing technologies gives rise to the abundance of large single-cell RNA-seq datasets at the scale of hundreds of thousands of single cells. Integrative analysis of large-scale scRNA-seq datasets has the potential of revealing *de novo* cell types as well as aggregating biological information. However, most existing methods fail to integrate multiple large-scale scRNA-seq datasets in a computational and memory efficient way. We hereby propose OCAT, **O**ne **C**ell **A**t a **T**ime, a graph-based method that sparsely encodes single-cell gene expressions to integrate data from multiple sources without most variable gene selection or explicit batch effect correction. We demonstrate that OCAT efficiently integrates multiple scRNA-seq datasets and achieves the state-of-the-art performance in cell type clustering, especially in challenging scenarios of non-overlapping cell types. In addition, OCAT efficaciously facilitates a variety of downstream analyses, such as differential gene analysis, trajectory inference, pseudotime inference and cell inference. OCAT is a unifying tool to simplify and expedite the analysis of large-scale scRNA-seq data from heterogeneous sources.

## 2 Introduction

The rapid advancement of transcriptome sequencing technologies in single cells (scRNA-seq) has witnessed the exponential growth in the number of large-scale scRNA-seq datasets. Integration of multiple scRNA-seq datasets from different studies has the great potential to facilitate the identification of both common and rare cell types, as well as *de novo* cell groups. Data heterogeneity, or batch effect, is one of the biggest challenges when integrating multiple scRNA-seq datasets. Batch effect is the perturbation in measured gene expressions, often introduced by factors such as library preparation, sequencing technologies and sample origins (donors). Batch effect is therefore likely to confound with true biological signals representing cell identities, resulting in misclassification of cells by experiment rather than by their true biological identities. Batch effect removal has thus become a mandatory step prior to data integration, introducing additional computational challenges. Most existing batch effect removal procedures assume that the biological effect is orthogonal to the batch effect, which is unlikely to be true in real life. Moreover, as the scale of the datasets increases, integrating multiple large-scale scRNA-seq datasets can induce heavy, or sometimes unbearable, computational and memory storage burden.

Most of the existing scRNA-seq integration methods require explicit batch removal steps. One of the most commonly used approach is mutual nearest neighbors (MNNs) [Haghverdi et al., 2018], which requires paired cells (or MNNs) to align the datasets into a shared space. However, this approach demands for large run-time memory and long computation time to search for MNNs in the high dimensional space of gene expressions. Though some derivatives of the MNN method [Hie et al., 2019, Polański et al., 2020] attempted to improve the memory efficiency by performing dimension reduction in the gene expression space, the memory usage is still demanding when the number of single cells is large. Another common approach, Seurat v3 [Stuart et al., 2019], projects scRNA-seq data to a canonical correlation analysis (CCA) subspace, and then computes MNNs in the CCA subspace to correct the batch effect. On the other hand, Harmony [Korsunsky et al., 2019] iteratively removes batch effects after projecting scRNA-seq data to a principal component analysis (PCA) subspace. However, Harmony can also consume large memory when the sample size is large. To reduce the computational burden of batch effect correction on scRNA-seq integration, we hereby propose OCAT (**O**ne **C**ell **A**t a **T**ime), a fast and memory-efficient machine learning-based method that does not require explicit batch effect removal in integrating multiple scRNA-seq datasets. OCAT utilizes sparse encoding to integrate multiple heterogeneous scRNA-seq datasets, achieving state-of-the-art or comparable performance compared to existing methods.

OCAT offers three major advantages over existing methods. First, OCAT identifies hypothetical “ghost” cells of each datasets and constructs a sparse bipartite graph between each cell with the “ghost” cells, generating a sparsified encoding of each cell optimized for computational efficiency (*O*(*N*)). Second, by connecting each individual cell to the “ghost” cell collection from all datasets, OCAT manages to capture the global similarity structure between single cells, and thus does not require any batch removal step. Thirdly, the OCAT sparse graph encoding can be effectively transformed into cell feature representations that readily tackle a wide range of downstream analysis tasks, providing a unified solution to common single-cell problems such as differential gene analysis, trajectory inference, psuedotime inference and cell inference.

## 3 Results

### 3.1 The OCAT framework overview

OCAT integrates multiple large-scale scRNA-seq datasets using sparse encoding as the latent representation of the single-cell gene expressions. Given multiple scRNA-seq gene expression matrices as input, OCAT first identifies hypothetical “ghost” cells, the centers of local neighbourhoods, from each dataset. OCAT then constructs a bipartite graph between all single cells to the “ghost” cell set using similarities as edge weights. OCAT further amplifies the strong connections and trims down the weak edges with the “ghost” cell set, by staying connected to the *s* most similar “ghost” cells only. We employ the Local Anchor Embedding (LAE) algorithm [Liu et al., 2010] to further optimize the edge weights from each single cell to the remaining *s* “ghost” cells, such that they can most effectively reconstruct the gene expression features of the single cell. These weights are then treated as the OCAT sparsified encoding of each single cell.

OCAT lastly captures the global cell-to-cell similarities through message passing between the “ghost” cells, which maps the sparsified weights of all single cells to the same global latent space. The sparsified weights are treated as the sparse encoding of each single cell.

As the number of the most similar “ghost” cells *s* is much smaller than the number of genes, the OCAT latent representation is very sparse. We show that this sparse encoding can effectively facilitate downstream analyses, such as cell type clustering, differential gene analysis, trajectory and pseudotime inference, as well as cell type inference. Moreover, OCAT sparse encoding is also capable of clustering spatial transcriptomics. Figure 1 outlines the workflow of the OCAT integration procedures as well as various downstream analysis functionalities.

**Figure 1:**
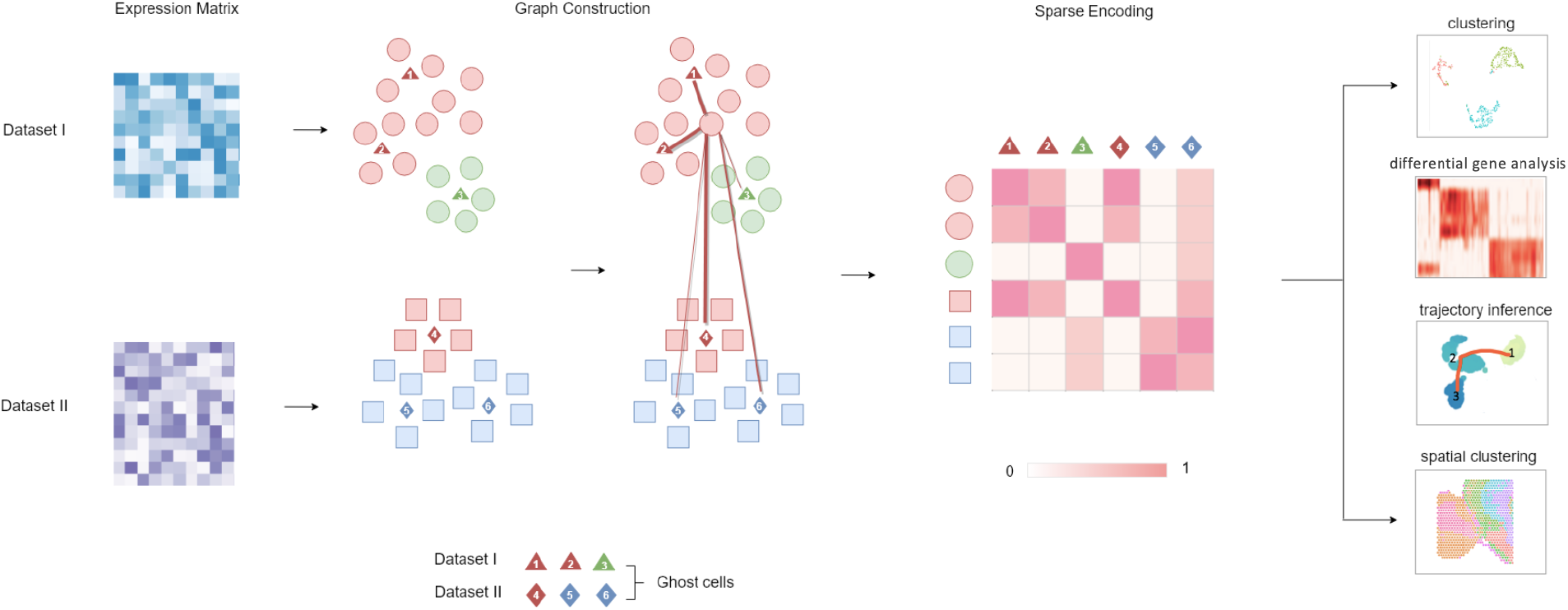
Schematic workflow of OCAT. For each individual scRNA-seq dataset, OCAT identifies “ghost” cells in each dataset, and encodes the gene expression of all single cells by their similarities to the “ghost” cell collection. This sparse encoding can effectively facilitate various downstream analysis tasks, such as cell clustering, differential gene analysis, trajectory inference and spatial scRNA-seq clustering.

### 3.2 Sparse encoding of single-cell transcriptomics effectively corrects batch effect and integrates multiple scRNA-seq datasets

When integrating multiple heterogeneous scRNA-seq datasets, most existing integration methods require iterations of explicit batch effect correction steps between every pair of datasets. Another common assumption in scRNA-seq data integration is that cell types are shared across all datasets, which is rarely true in real life. Such requirements and assumptions pose major challenges in the performance as well as computational efficiency to most existing integration methods. OCAT captures the global cell-to-cell similarity across datasets by connecting each single cell to the “ghost” cell set. OCAT thus does not require any explicit batch effect correction step and proves to be robust in identifying non-overlapping cell types unique to some datasets. The sparsified encoding also greatly accelerates the computational speed and considerably reduces the memory usage when integrating multiple large-scale datasets.

One common assumption of existing integration methods is that any cell type present in one dataset must also be present in all the other datasets. However, if some non-overlapping cell types exist, such methods can falsely cluster cells of the non-overlapping cell types. We here demonstrate how this assumption introduces misclassification in the presence of non-overlapping cell types. The human dendritic dataset [Villani et al., 2017] consists of human blood dendritic cell (DC), namely, CD1C DC, CD141 DC, plasmacytoid DC (pDC), and double negative cells. Tran et al. [2020] further processed and manually split the data into two batches: batch 1 contains 96 pDC, 96 double negative and 96 CD141 cells, while batch 2 has 96 pDC, 96 double negative and 96 CD1C cells. CD141 cells are only present in batch 1, while CD1C cells are only present in batch 2. The visualization of cell type clustering in Figure 2A shows that Seurat v3, Harmony and Scanorama all falsely group CD141 and CD1C together. On the other hand, OCAT manages to distinguish CD141 and CD1C as two separate cell clusters. This verifies that by constructing the single cell to “ghost” cell bipartite graph, the OCAT sparse encoding successfully recovers global cell-to-cell similarity across batches and captures true cell type identities. The cell type clustering metrics also reflect the same result, where OCAT has NMI_cell type_ = 0.7718, higher than all the other benchmarked methods; see Table 1 for a detailed comparison.

**Figure 2:**
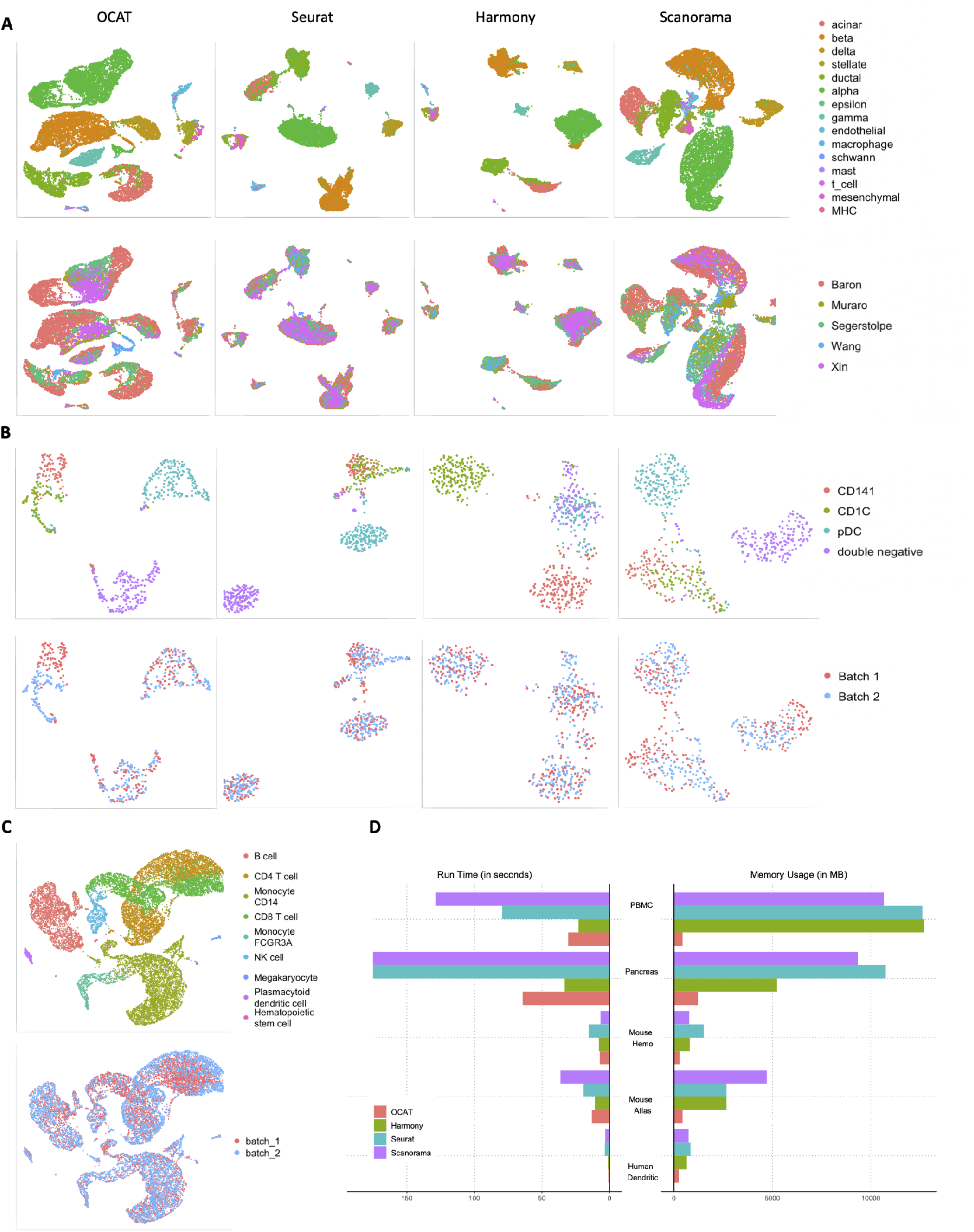
Integrating multiple scRNA-seq datasets with OCAT. **A**: UMAP projection of five integrated human pancreatic scRNA-seq datasets from heterogeneous sequencing platforms by OCAT, Seurat v3, Harmony, and Scanorama. The top panel is colored by the annotated cell types and the bottom panel is colored by the dataset. **B**: UMAP projection of two integrated human dendritic datasets with non-overlapping cell types by OCAT, Seurat v3, Harmony, and Scanorama. The top panel is colored by the annotated cell types and the bottom panel is colored by the dataset. **C**: UMAP projection of two integrated PBMC scRNA-seq datasets by OCAT; see Supplementary Figure S1 for a full comparison with Seurat v3, Harmony and Scanorama. **D**: Memory usage and runtime of OCAT, Seurat v3, Harmony and Scanorama on five scRNA-seq integration tasks.

**Table 1:** Cell type clustering and batch correction performance on integrating multiple scRNA-seq datasets. The clustering performance is measured by the Normalized Mutual Information (NMI), where NMI_cluster_ = 1 implies correct clustering by cell type annotations. (1 − NMI_batch_) = 1 implies no batch effect present after integration; see Supplementary Material 2.2.2-2.2.3 for more details on NMI_cluster_ and NMI_batch_, and Supplementary Table S1 for additional evaluation metrics.

| Dataset | $n_{\text{cells}}$ | $n_{\text{genes}}$ | Metric | OCAT | Seurat | Harmony | Scanorama |
| --- | --- | --- | --- | --- | --- | --- | --- |
| Human dendritic | 576 | 16,594 | $\text{NMI}_{\text{cluster}}$ | **0.7718** | 0.7375 | 0.7653 | 0.7212 |
| | | | $1 - \text{NMI}_{\text{batch}}$ | 0.9999 | 0.9999 | **1.0000** | 0.9999 |
| Mouse atlas | 6,954 | 5,558 | $\text{NMI}_{\text{cluster}}$ | **0.8006** | 0.6981 | 0.7625 | 0.6960 |
| | | | $1 - \text{NMI}_{\text{batch}}$ | **0.9999** | 0.9970 | 0.9959 | 0.9905 |
| Human pancreas | 14,767 | 15,558 | $\text{NMI}_{\text{cluster}}$ | **0.7949** | 0.7947 | 0.7302 | 0.7249 |
| | | | $1 - \text{NMI}_{\text{batch}}$ | 0.9650 | **0.9762** | 0.9600 | 0.9538 |
| PBMC | 15,476 | 17,430 | $\text{NMI}_{\text{cluster}}$ | 0.7424 | **0.7932** | 0.7851 | 0.7141 |
| | | | $1 - \text{NMI}_{\text{batch}}$ | 0.9934 | 0.9943 | **0.9944** | 0.9939 |
| Mouse hematopoietic | 4,649 | 3,467 | $\text{NMI}_{\text{cluster}}$ | 0.5019 | 0.4606 | 0.4111 | **0.5160** |
| | | | $1 - \text{NMI}_{\text{batch}}$ | 0.9648 | 0.9673 | **0.9734** | 0.9674 |

We then demonstrate the performance and efficiency of OCAT on integrating more than two heterogeneous scRNA-seq datasets. The pancreatic dataset consists of five human pancreatic scRNA-seq datasets sequenced with four different technologies (inDrop [Baron et al., 2016], CEL-Seq2 [Muraro et al., 2016], Smart-Seq2 [Segerstolpe et al., 2016], SMARTer [Wang et al., 2016, Xin et al., 2016]). Datasets generated by different sequencing platforms and technologies have inherent technical differences [Hicks et al., 2018, Tung et al., 2017], posing greater challenge to the integration task as the distributions of gene expressions vary significantly across the five datasets. Another challenge lies in the computational cost and memory consumption of integrating five datasets, caused by the iterative batch correction process for large number of cells with high dimensional gene expressions. Nevertheless, OCAT outperforms the other methods in correctly identifying the cell types without any most variable gene selection or batch removal steps. Following the data pre-processing procedures outlined in Tran et al. [2020], we integrate the five pancreatic datasets using OCAT, and benchmark with three existing integration methods, Seurat v3 [Stuart et al., 2019], Harmony [Korsunsky et al., 2019] and Scanorama [Hie et al., 2019]. The UMAP projection in Figure 2A demonstrates that OCAT outperforms the other methods in identifying the cell types (NMI_cell type_ = 0.8037), while achieving comparable batch correction performance (1 − NMI_batch_ = 0.9638); see Table 1 for details. We show in Figure 2D that OCAT is more computationally and memory efficient than the other benchmarked methods. Notably, OCAT takes less than half of the runtime of Seurat v3 and Scanorama. Though Harmony runs slightly faster than OCAT, it consumes four times more memory than OCAT. Seurat v3 and Scanorama both require more than 8 times memory of OCAT.

We also validate the performance of OCAT on integrating mouse cell atlas [Han et al., 2018], human peripheral blood mononuclear cell (PBMC) [Xin et al., 2016] and mouse hematopoietic stem and progenitor cell [Nestorowa et al., 2016] datasets. OCAT achieves state-of-the-art or comparable performance with the other benchmarked methods; see Table 1, Figure 2C and Supplementary Figure S1-3 for details. Notably, when integrating the two PBMC datasets with a total of 15,476 single cells and 33,694 genes, OCAT is twice faster than Seurat v3 and three times faster than Scanorama. In addition, Harmony and Seurat v3 consume more than 29 times of OCAT’s memory usage, while Scanorama consumes more than 24 times of OCAT’s memory usage (Figure 2D).

### 3.3 OCAT unifies various downstream biological inferences

The OCAT sparse encoding framework can readily extract the latent representations of single cells from individual scRNA-seq datasets. We show in this section that the OCAT sparse encoding can effectively facilitate various downstream analyses with important biological implications, such as cell inference, differential gene analysis, as well as trajectory inference and psuedotime inference. We further show that OCAT can also extract sparse latent representation of spatial scRNA-seq datasets.

#### 3.3.1 Sparse encoding of individual scRNA-seq dataset

We first demonstrate with four large-scale scRNA-seq datasets, Romanov [Romanov et al., 2017], Zeisel [Zeisel et al., 2015], Retina [Shekhar et al., 2016], and PBMC 68k [Zheng et al., 2017]. We show in Table 2 that OCAT consistently outperforms existing methods in cell type clustering [Wang et al., 2017, Stuart et al., 2019, Lopez et al., 2018]; see Figure 3 for the UMAP visualizations. OCAT is also capable of sparsely encoding spatial scRNA-seq data by treating the spatial coordinates as additional feature representations of single cells. We show in Figure 3E the identification of cell types using the OCAT sparse encoding on a sagital mouse brain spatial scRNA-seq dataset [Stuart et al., 2019].

**Figure 3:**
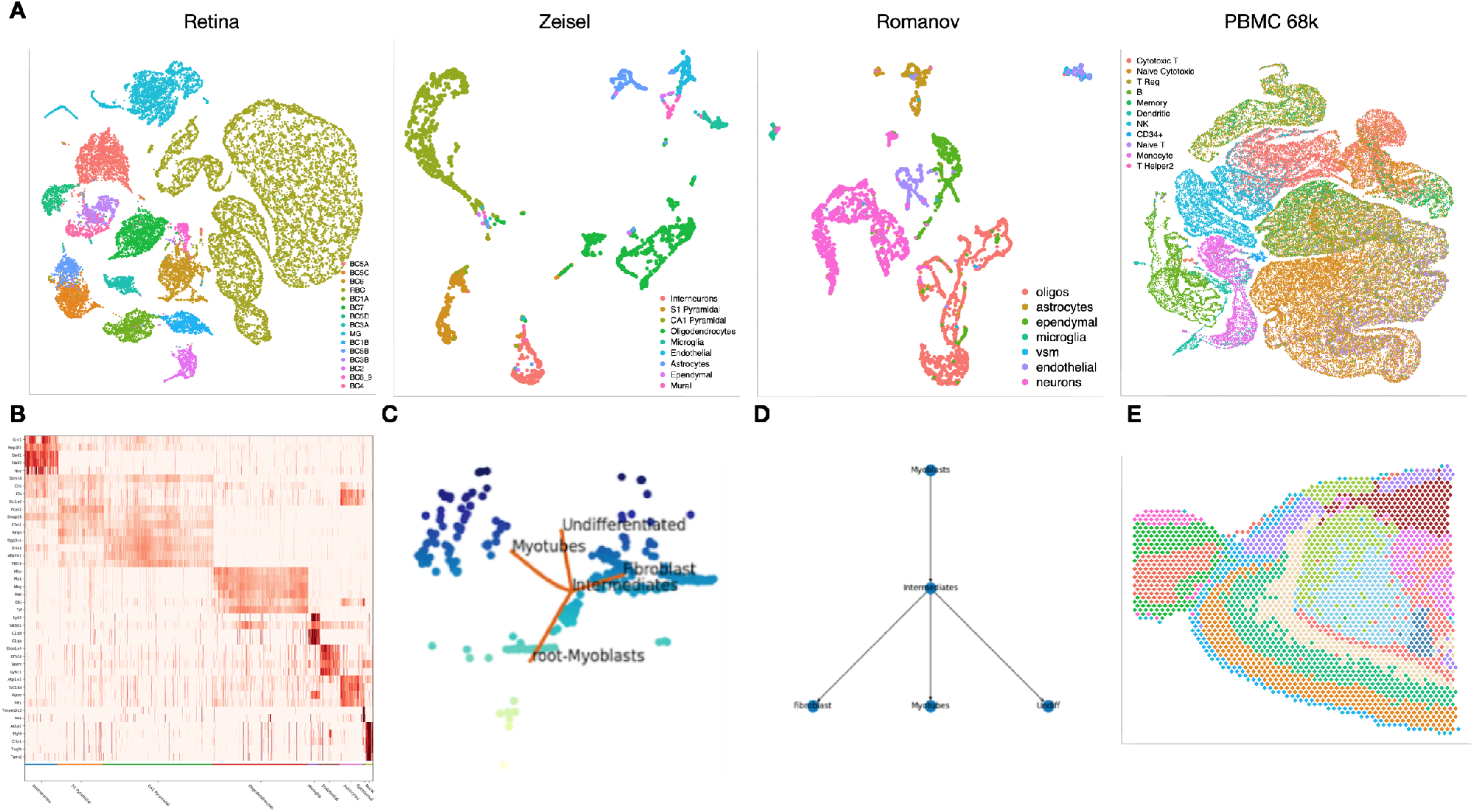
OCAT on individual scRNA-seq datasets. **A**: UMAP projection of the OCAT sparsified embeddings for Romanov, Zeisel, Retina and PBMC 68k datasets, colored by the annotated cell types. **B**: Differential gene analysis with OCAT on the Zeisel dataset. **C-D**: OCAT trajectory inference and pseudotime inference on the HSMM dataset. **E**: Spatial scRNA-seq clustering using the OCAT sparsified embeddings.

**Table 2:** **Clustering performance of OCAT** on four individual scRNA-seq datasets: Romanov, Zeisel, Retina and PBMC 68k, benchmarked with scVI, Seurat v3 and SIMLR. The clustering performance is measured by the Normalized Mutual Information (NMI), where NMI = 1 implies correctly clustering all the cells with the same cell types while NMI = 0 indicates random guessing; see Supplementary Table S5 for additional evaluation metrics.

| Dataset | $n_{\text{cells}}$ | $n_{\text{genes}}$ | OCAT | scVI | Seurat | SIMLR |
| --- | --- | --- | --- | --- | --- | --- |
| Romanov | 2,881 | 24,341 | **0.6443** | 0.5436 | 0.6343 | 0.4234 |
| Zeisel | 3,005 | 19,972 | **0.7884** | 0.7126 | 0.6724 | 0.7373 |
| Retina | 19,829 | 13,166 | **0.8742** | 0.7572 | 0.7865 | 0.6337 |
| PBMC 68k | 68,579 | 1,000 | **0.5750** | 0.4638 | 0.4899 | 0.5344 |

#### 3.3.2 Cell type inference

OCAT supports immediate and accurate cell type inference of incoming data, without repeating feature extraction procedures combining the incoming data with the existing database. We denote the existing scRNA-seq datasets as the “reference” dataset, and the incoming unlabelled data as the “inference” dataset. For the reference dataset, OCAT first identifies a set of “ghost” cells and extracts the sparse features to train a Support Vector Machine (SVM) [Noble, 2006] with the annotated cell types. For the inference cells, OCAT computes their sparse encoding using the pre-identified reference “ghost” cell set and transfers cell type labels to the incoming inference cells using the pre-trained SVM.

We first demonstrate OCAT’s cell type inference performance on four individual scRNA-seq datasets, Romanov [Romanov et al., 2017], Zeisel [Zeisel et al., 2015], Retina [Shekhar et al., 2016] and PBMC 68k [Zheng et al., 2017]. Each dataset is randomly split into 90% reference set and 10% inference set. The OCAT-extracted features of the inference cells based on the reference “ghost” cells yield high accuracy in cell type assignment; see Supplementary Table S5 and Supplementary Figure S5. We further demonstrate that OCAT can infer cell types in a more challenging scenario across two heterogeneous scRNA-seq datasets. With two PMBC scRNA-seq datasets [Polański et al., 2020], we split each dataset into 90% reference set and 10% inference set. OCAT assigns cell types to the 10% inference set from dataset 2 based on the 90% reference set from dataset 1, and vice versa, achieving F1 score of 0.8907 and 0.7719 respectively. We also conducted cell inference experiments on two mouse atlas datasets [Han et al., 2018, Consortium et al., 2018] and two human dendritic datasets [Villani et al., 2017, Tran et al., 2020], both achieving high accuracy in cell type assignment; see Supplementary Figure S5 for details.

#### 3.3.3 Differential gene analysis

Differential gene analysis is one of the most common approaches to facilitate cell type annotations. OCAT effectively selects the marker genes for each cell group based on the raw gene expression data. We demonstrate the efficacy of OCAT in differential gene analysis using the Zeisel dataset [Zeisel et al., 2015] that classifies 9 cell types in the mouse somatosensory cortex and hippocampal CA1 region. Figure 3B plots the top 5 marker genes for each cell type. OCAT manages to replicate the marker gene findings reported by [Zeisel et al., 2015], for example, *Gadl* and *Gad2* genes for Interneuron cells, and *Acta2* gene for mural cells. We further compare the top selected differential genes by OCAT with those identified by Seurat v3, and show that the top selected genes are highly consistent between the two methods. For example, for the CA1 cell population, OCAT identifies *Crym, Cpne6, Neurod6, Gria1* and *Wipf3* as the top five differential genes, and four of them are also in the top five differential genes selected by Seurat v3. We further show that the top features are highly consistent for all other cell populations in Supplementary Table S6 and Supplementary Figure S6.

#### 3.3.4 Trajectory and pseudotime inference

OCAT is able to reconstruct the developmental trajectory and pseudotime of cells from their transcriptomic profiles. In most cell populations, there exists a gradient of differentiation underlying the process of cell renewal, from progenitor cells to the terminally differentiated cell types. Based on the similarities in gene expressions, trajectory and pseudotime analyses infer the differentiation status of the cell types as well as individual cells. Trajectory inference first maps out the developmental lineages from the least differentiated to most differentiated cell types. Pseudotime analysis then orders the individual cells along the predicted lineages and assigns each cell a pseudotime, indicating its time stamp in the process of differentiation.

OCAT extracts a reduced “ghost” neighbourhood graph between cell types by aggregating cell-to-cell similarities in each cluster. OCAT then infers the lineages by constructing the minimal spanning tree [Kruskal,1956] over the aggregated “ghost” neighbourhood graph that connects all the cell types. The least differentiated cell type is considered as the root cluster, which determines the unique directionality of the inferred lineages; see Section 5 for details. Lastly, to compute the pseudotime of each cell, OCAT appoints the least differentiated cell in each “ghost” neighbourhood as the root cell. Traversing down the lineages, OCAT uses the root cell as the point of reference in each local neighbourhood to assign pseudotime to individual cells.

We validate the performance of OCAT trajectory and pseudotime inference using the human skeletal muscle myoblast (HSMM) dataset [Trapnell et al., 2014]. The HSMM dataset contains time-series scRNA-seq data outlining the early stages of myogenesis. The 271 myoblast cells were collected at 0, 24, 48 and 72 hours of differentiation, with gold-standard annotations based on known gene markers [Tran and Bader, 2020, Trapnell et al., 2014]. OCAT infers the differentiation trajectory from myoblast to intermediate cells, followed by three separate branches into myotubes, fibroblasts and undifferentiated cells. Fibroblasts and undifferentiated cells represent the two cell groups that exit the differentiation cycle prior to myotube formation. The inferred trajectory is consistent with the known biology of myotube formation as well as the original findings in Tran and Bader [2020]. The pseudotime assigned by OCAT is highly correlated with the collection time stamps, with a Pearson correlation of 0.8743 by annotated cell type group. Additionally, following the procedures in Saelens et al. [2019], we compare OCAT with Slingshot [Street et al., 2018], PAGA Tree [Wolf et al.,2019] and Monocle ICA [Qiu et al., 2017] on trajectory and pseudotime inference with 28 gold-standard real datasets using the `dynverse R` package [Saelens et al., 2019]. OCAT is competitive in accurately assigning cell positions along the lineages as well as assisting downstream tasks of identifying important genes specific to the trajectory; see Supplementary Material 6.6.3, Supplementary Figure S7, and Supplementary Table S7 for details.

## 4 Discussion

In this work, we present OCAT, a unifying framework for analyzing large-scale scRNA-seq datasets, which synergizes a wide range of downstream tasks crucial to biological discoveries. OCAT utilizes sparse encoding as the latent representation of single cells to amplify the true biological signals. Through the hypothetical “ghost” cells, the OCAT sparse encoding captures the global cell-to-cell similarity across multiple datasets. We demonstrate that, without any batch effect correction, the sparse encoding of OCAT effectively separates the true biological differences among the cells from batch effects, achieving state-of-the-art performance with existing methods in cell type itentification.

Unlike most existing methods, OCAT does not rely on most variable gene selection to discriminate biological cell groups, which preserves the identities of non-overlapping cell types unqiue to some datasets and has the potential to facilitate the discovery of *de novo* cell groups. Furthermore, OCAT successfully leverages the high demand for computational resources in integrating large-scale scRNA-seq datasets. Through its sparse encoding of gene expressions, OCAT can scale up to integrate multiple scRNA-seq datasets with large number of cells and large number of genes, in a computational and memory efficient way.

OCAT effectively facilitates a variety of downstream analyses with important biological implications. For example, OCAT is readily applicable to analyzing individual scRNA-seq dataset as well as spatial scRNA-seq data, outperforming existing methods in cell type clustering. Moreover, OCAT can undertake challenging tasks such as differential gene analysis, trajectory inference, pseudotime inference and cell inference. With additional biological priors, OCAT has the great potential to better facilitate downstream analyses and extend to tackle more complex tasks such as cell-to-cell communication network inference, which we will explore as future work.

OCAT is freely available at https://github.com/bowang-lab/OCAT.

## 5 Online methods

### 5.1 The OCAT framework overview

OCAT endorses sparse encoding of the latent representations of the single-cell transcriptomics. Given *N* single cells each with *M* gene expressions, OCAT first identifies *m* ≪ *M* “ghost” cells and connects each individual cell with the ghost cells through a bipartite graph where the weights of the edges are treated as the encoding. As *m* ≪ *M*, the OCAT encoding is very sparse and computationally fast for large-scale datasets. The sparse encoding can then be deployed to find the similarities between cells within the same datasets as well as across multiple datasets. The similarities between cells can facilitate downstream analyses such as cell clustering, trajectory inference, gene prioritization. Moreover, the OCAT sparse encoding is also capable of clustering spatial transcriptomics. Figure 1 depicts the sparse encoding procedures of OCAT and the latent representations for various downstream tasks given two input scRNA-seq datasets. In the next sections, we will outline the OCAT algorithms in details.

### 5.2 Data pre-processing

Given an *N* × *M* gene expression matrix, *R*, where *N* is the number of single cells and *M* is the number of genes, OCAT first pre-processes the raw gene expression data by log-transforming each entry *r_ij_* in *R*

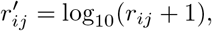

and normalizes the log-transformed expression to

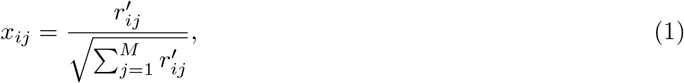

for *i* = 1, …, *N* and *j* = 1, …, *M*. The normalized gene expression matrix is denoted as 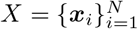, where ***x**_i_* = (*x*_*i*1_, *x*_*i*2_, … ,*x_iM_*)^*T*^ is the *M* × 1 normalized gene expression vector of cell *i*.

### 5.3 Dimension reduction of gene expression matrix

To efficiently encode the transcriptomics of the single cells, OCAT further reduces the dimension of the normalized gene expression matrix *X*. OCAT adopts the online Fast Similarity Matching (FSM) algorithm [Giovannucci et al., 2018] that projects each ***x**_i_* from 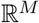 to 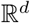 such that

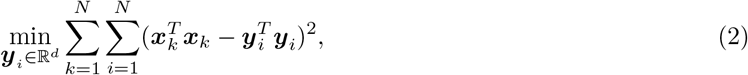

where ***y**_i_* is the *d* × 1 feature vector for cell *i*. OCAT adopts 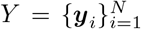 as the transcriptomic feature representation that will facilitate the construction of the sparsified bipartite graph.

Note that though a vast collection of methods is available for dimension reduction, online FSM is much more efficient with a complexity of *O*(*NMd*) than the traditional principal component analysis (PCA) whose complexity is *O*(*M*^2^*N* + *M*^3^), which is offered as an alternative option in the OCAT software package.

### 5.4 Sparsified bipartite graph for single-cell transcriptomics

#### 5.4.1 Identifying “ghost” cells

We introduce the idea of “ghost” cells which are imaginary cells that characterize the transcriptomics of the real single cells. OCAT identifies *m* ghost cells that are the K-Means cluster centers of *Y*, and denotes their features as ***u**_j_*, for *j* = 1, 2, …, *m*. We then construct a sparsified bipartite graph *G* = (*V, U, E*) between the single cells and the ghost cells, where each node *v_i_* represents the feature ***y**_i_* of the ith single cell.

#### 5.4.2 Construct sparsified bipartite graph

Our next goal is to construct the sparsified bipartite graph between the single cells and the ghost cells. For single cell *i*, the weights ***z**_i_* = (*Z*_*i*1_, *Z*_*i*2_, …, *Z_im_*)^*T*^ is a *m* × 1 vector such that

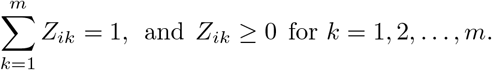

For single cell *i*, OCAT first identifies its *s* closest ghost cells with the top *s* cosine similarity values and denote their indices as 〈*i*〉 ∈ [1 : *m*]. OCAT then optimizes the edge weights cell *i* and its *s* neighbor ghost cells, ***z***_〈*i*〉_, using Local Anchor Embedding (LAE) [Liu et al., 2010] by

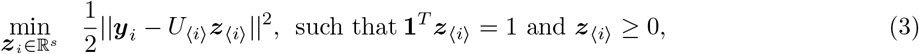

and *U*_〈*i*〉_ = {***u**_k_*}_*k*∈〈*i*〉_ are the features of the *s* neighbor ghost cells. The edge weights of cell *i* to all the ghost cells are thus denoted as as 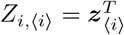 and 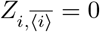, and the collection of all the edge weights connecting *N* single cells to *m* ghost cells is denoted as 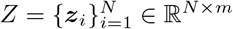.

#### 5.4.3 Message passing between single cells

To infer the transcriptomic similarity between single cells, a common approach is to compute the adjacency matrix *W* between the cells. However, when the number of single cells, *N*, is large, storing a *N* × *N* adjacency matrix consumes significant memory. OCAT, instead of computing cell-to-cell similarity directly, infers it through single cell to ghost cell edge weights, *Z*, and the similarities between ghost cells, 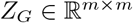. The similarity between ghost cells is defined as,

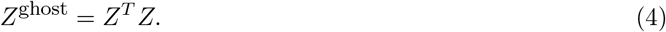

We then standardize *Z*^ghost^ by

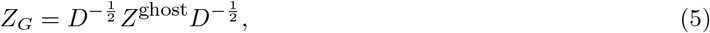

where *D* is a diagonal matrix with

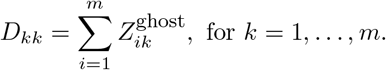

The normalized ghost cell to ghost cell similarity, *Z_G_*, is an *m* × *m* combined ghost cell set that transmits messages between single cells. Lastly, we obtain refined sparse embeddings for the single cells through message passing *Z_W_* by

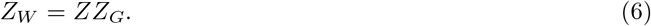

### 5.5 Integration of multiple scRNA-seq datasets

OCAT can easily integrate multiple gene expression datasets thanks to the design of sparsified bipartite graph. Without loss of generality, suppose we have two scRNA-seq datasets to integrate, each with *N*_1_ and *N*_2_ single cells and *M* common genes. Each individual dataset first undergoes the same pre-processing and dimension reduction steps outlined in Section 5.2 and 5.3, and yields 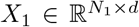 for dataset 1 and 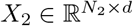 for dataset 2.

OCAT then identifies *m*_1_ ghost cells from *X*_1_ with features 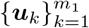, and *m*_2_ ghost cells from *X*_2_ with features 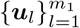. For the ith individual cell, OCAT identifies *s*_1_ closest ghost cells with indices 〈*i*1〉 from the first ghost cell set and *s*_2_ closest ghost cells with indices 〈*i*2〉 from the second set. Within the first ghost cell set, OCAT obtains the optimized weights ***z***_〈*i*1〉_ such that

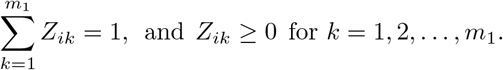

Similarly, the optimized weights for the second ghost cell set, OCAT obtains the optimized weights ***z***_〈*i*2〉_. The weights of the edges connecting the ith single cell to all the ghost cells are then denoted as *Z*_i,〈*i*〉_ = (***z***_〈*i*1〉_, ***z***_〈*i*2〉_)^*T*^ and 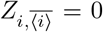. The collection of all the edge weights of (*N*_1_ + *N*_2_) single cells connecting to (*m*_1_ + *m*_2_) ghost cells is denoted as 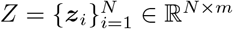 where *N* = *N*_1_ + *N*_2_ and *m* = *m*_1_ + *m*_2_.

Following (4) and (5), we obtain the re-fined embeddings, 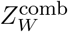, for each single cell through message passing between the combined ghost cells. We lastly normalize 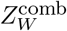 by

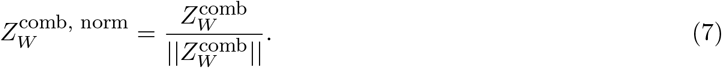

### 5.6 Differential gene analysis

OCAT offers the functionality to find the differential genes for each cell type clusters. Denote the normalized gene expression matrix as *X* = {*x_ij_*}, where *i* = 1, …, *N* and *j* = 1, …, *M*. For cell type cluster *C*, we compares the gene expression of cell type cluster *C* with all the other types, and we rank the top differential genes by the magnitude of

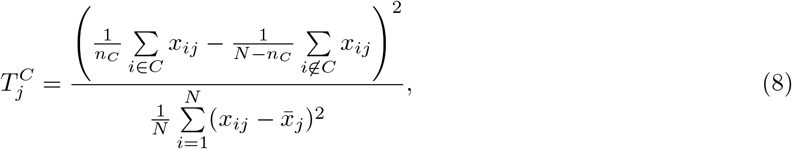

where 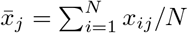.

### 5.7 Cell inference

OCAT supports immediate cell type inference of incoming data based on existing databases, without recomputing the latent representations by combining the new incoming (“inference”) dataset and the existing (“reference”) dataset.

Given an incoming “inference” set, OCAT first projects the normalized gene expression *X*^infer^ to the same 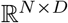 subspace as the “reference” set, obtaining the reduced cell representation *Y*^infer^. OCAT then constructs a bipartite graph that connects these new “inference” cells to the “ghost” cells identified in the “reference” set following (3), and obtains the edge weights, *Z*^infer^, for the “inference” cells. The edge weights then go through the same message-passing procedures as the “reference” cells, resulting in 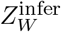, the sparse representation of the new “inference” cells mapped to the same global subspace as the “reference” cells.

To assign cell type labels to the “inference” cells, OCAT “trains” a Support Vector Machine (SVM) [Noble, 2006] based on the sparse representations of the “reference” cells, 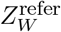, and the cell type labels for the “reference” cells. Based on the estimated coefficients from SVM, OCAT infers the cell type labels of the new incoming cells using 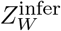.

### 5.8 Trajectory inference

Trajectory inference aims to computationally reconstruct the developmental trajectory of cells based on gene expressions. It outlines the temporal transition from the the least differentiated to the most differentiated cell types. OCAT infers the developmental lineages by connecting the similarity graph between cell types with a minimum spanning tree [Kruskal, 1956].

Suppose we have an *N* × *m* dimensional gene expression embedding for the cells, for example, the sparse embedding by OCAT, 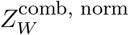. The cells are clustered into *c* cell types based on the embedding. OCAT computes the similarity score between cell type *p* and cell type *q*, *A_p,q_*, by averaging the pair-wise cell-to-cell cosine similarities between cell types *p* and *q*.

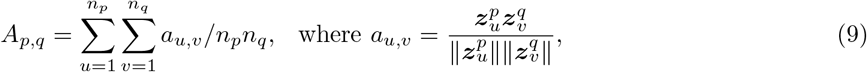

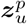 is the embedding vector for the *u*th cell in cell type *p*, 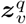 is the embedding vector for the *v*th cell in cell type *q*, and *n_p_*, *n_q_* are the number of cells in cell type *p* and *q*, respectively. ∥ · ∥ denotes the l2-norm.

Let 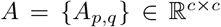 denote the matrix of pair-wise similarity scores between *c* cell types. OCAT constructs an undirected graph *G^C^* from *A*, where each node represents a unique cell type, and the edge weight between two nodes (two cell types) is their similarity score. OCAT then obtains the minimum spanning tree *T* that connects all the nodes while minimizing the total sum of edge weights in the tree *T*. OCAT lastly adds directionality to the tree by taking the least differentiated cell type, namely, the root cell type, as the starting point of differentiation. Once the root cell type is determined, we obtain a unique directionality within the tree *T*.

### 5.9 Pseudotime Inference

Pseudotime analysis assigns each cell a time stamp along the lineages: less differentiated cells have earlier time stamps; more differentiated cells have later time stamps. It thus provides more granularity to individual cells than the lineage ordering of cell types. OCAT defines a root cell in the root cluster, ***r***_1_, to serve as a reference to quantify differentiation. Biologically, ***r***_1_ represents the most primitive in the entire differentiation trajectory. OCAT identifies ***r***_1_ computationally by locating the cell whose spatial distances with other cells have the best accordance with the lineage ordering of cell types identified. OCAT then infers the extent to which a particular cell differentiates using its distance to the most primitive ***r***_1_, where less differentiated cells are closer to ***r***_1_, and vice versa.

To calculate the distance of the *u*th cell of type *p* to the first cell ***r***_1_, OCAT considers both the position of cluster *p* along the cell type lineages and the position of cell *u* in cluster *p*. We then define a root cell in every non-root cluster to serve as landmarks to connect the cell types along the lineages, denoted as 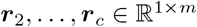. In a non-root cluster *p*, the cell with the closest average Euclidean distance with all cells in the previous cluster *p* − 1 on the same lineage is assigned to be the root cell, ***r**_p_*. OCAT defines a distance *D_i_* for each cell in the dataset, where 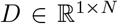. The distance for the *u*th cell in cluster *p* is defined as the sum of Euclidean distance between 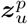 and the current root cell cluster ***r**_p_*, and the length of cell type lineages up to cluster *p*:

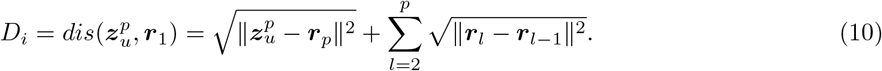

OCAT uses the normalized distance Dnorm as the pseudotime measure:

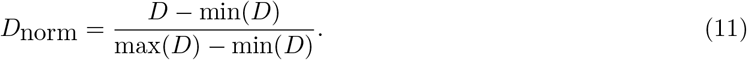

