## Supplementary Materials for "One Cell At a Time: A Unified Framework to Integrate and Analyze Single-cell RNA-seq Data"

### One Cell At a Time – Supplementary Material

July 8, 2021

**7 Data and code availability** **14**

### 1 Additional methods

#### 1.1 Efficient dimension reduction of the gene expression matrix

In the dimension reduction step, OCAT adopts the online Fast Similarity Matching (FSM) [4] to efficiently project the normalized gene expression $X \in \mathbb{R}^M$ to its principal subspace $Y \in \mathbb{R}^d$. The online FSM algorithm is a fast and memory-efficient method for principal subspace projection (PSP), which outputs an updated estimate of the principal subspace after intaking a new datum (one cell at a time). The online FSM is an improvement over the Similar Matching (SM) [14] algorithm that solves

$$\min_{W} \max_{M} \; 2\mathrm{Tr}(W^T W) - \mathrm{Tr}(M^T M) - 2\sum_{t=1}^{N} \boldsymbol{x}_t^T W^T \boldsymbol{y}_t, \tag{1}$$

where $\boldsymbol{y}_t \equiv M^{-1} W \boldsymbol{x}_t$, for $t = 1, 2, \ldots, N$. As outlined in Algorithm 1, SM costs $O(Md + d^3)$ per iteration to solve

$$M\boldsymbol{y}_t = W\boldsymbol{x}_t. \tag{2}$$

When the reduced dimension $d \sim \sqrt{M}$, the cost $O(d^3)$ for solving is non-negligible. To accelerate the SM algorithm, the online FSM algorithm (Algorithm 2) adopts the Sherman–Morrison formula when updating $M_{\mathrm{inv}}$, thus reducing the computing cost to $O(Md)$ per iteration.

**Algorithm 1** Similarity Matching (SM) [14, 15]

**Input:** Initial weights $M \in \mathbb{R}^{K\times K}$ and $W \in \mathbb{R}^{K\times D}$

**for** $t = 1, 2, 3, \ldots$ **do**
  $\boldsymbol{y}_t \leftarrow M^{-1} W \boldsymbol{x}_t$
  $W \leftarrow (1-\alpha_t)W + \alpha_t \boldsymbol{y}_t \boldsymbol{x}_t^T$
  $M \leftarrow (1-\beta_t)M + \beta_t \boldsymbol{y}_t \boldsymbol{y}_t^T$
**end for**

**Algorithm 2** Fast Similarity Matching (FSM) [4]

**Input:** Initial weights $M_{\mathrm{inv}} \in \mathbb{R}^{K\times K}$ and $W \in \mathbb{R}^{K\times D}$

**for** $t = 1, 2, 3, \ldots$ **do**
  $\boldsymbol{y}_t \leftarrow M_{\mathrm{inv}}^{-1} W \boldsymbol{x}_t$
  $M_{\mathrm{inv}} \leftarrow \frac{1}{1-\beta_t} M_{\mathrm{inv}}$
  $\boldsymbol{z}_t \leftarrow M_{\mathrm{inv}} \boldsymbol{y}_t$
  $M_{\mathrm{inv}} \leftarrow M_{\mathrm{inv}} - \frac{\beta_t}{1+\beta_t \boldsymbol{z}_t^T \boldsymbol{y}_t} \boldsymbol{z}_t \boldsymbol{z}_t^T$
**end for**

#### 1.2 Sparse graph construction through latent anchor embedding

OCAT constructs a sparse bipartite graph that connects each single cell to the "ghost" cell set. To compute the edge weights, $\boldsymbol{z}_{\langle i \rangle}$, between cell $i$ and its $s$ closest "ghost" cells, OCAT adopts the Local Anchor Embedding (LAE) algorithm [8] to optimize

$$\min_{\boldsymbol{z}_i \in \mathbb{R}^s} \; \frac{1}{2}||\boldsymbol{y}_i - U_{\langle i \rangle}\boldsymbol{z}_{\langle i \rangle}||^2, \text{ such that } \mathbf{1}^T \boldsymbol{z}_{\langle i \rangle} = 1 \text{ and } \boldsymbol{z}_{\langle i \rangle} \geq 0, \tag{3}$$

and $U_{\langle i \rangle} = \{\boldsymbol{u}_k\}_{k \in \langle i \rangle}$ are the features of the $s$ neighbor ghost cells.

LAE applies the projected gradient method to solve (3), and uses the Nesterov's method [11] to accelerate the gradient decent step (Algorithm 3). The LAE algorithm outputs a highly sparse weight matrix $Z$, with a memory usage of $O(sN)$ and time complexity $O(smN + s^2TN)$, where $m$ is the total number of candidate "ghost" cells, $s$ is the number of closest "ghost" cells to be selected, and $T$ is the number of iterations. In practice, the LAE algorithm converges within a few iterations and $T$ is therefore small. The LAE algorithm significantly accelerates the computational efficiency and reduces the memory usage of OCAT.

**Algorithm 3** Local Anchor Embedding (LAE)

**Input:** data points $\{\boldsymbol{x}_i\}_{i=1}^n \subset \mathbb{R}^d$, anchor point matrix $U \in \mathbb{R}^{d\times m}$, integer s.

**for** $i$ **to** $n$ **do**

for $\boldsymbol{x}_i$ find $s$ nearest neighbors in $U$, saving the index set $\langle i \rangle$;

define functions $g(z) = ||\boldsymbol{x}_i - U_{\langle i \rangle}\boldsymbol{z}||^2/2$, $\nabla g(\boldsymbol{z}) = U_{\langle i \rangle}^T U_{\langle i \rangle}\boldsymbol{z} - U_{\langle i \rangle}^T \boldsymbol{x}_i$, and $\tilde{g}_{\beta,\boldsymbol{v}}(\boldsymbol{z}) = g(\boldsymbol{v}) + \nabla g(\boldsymbol{v})^T(\boldsymbol{z} - \boldsymbol{v}) + \beta||\boldsymbol{z} - \boldsymbol{v}||^2/2$;

initialize $\boldsymbol{z}^{(0)} = \boldsymbol{z}^{(1)} = \mathbf{1}/s$, $\delta_{-1} = 0$, $\delta_0 = 1$, $\beta_0 = 1$, $t = 0$;

**repeat**

$t = t + 1$, $\alpha_t = \frac{\delta_{t-2}-1}{\delta_{t-1}}$

set $\boldsymbol{v}^{(t)} = \boldsymbol{z}^{(t)} + \alpha_t(\boldsymbol{z}^{(t)} - \boldsymbol{z}^{(t-1)})$

**for** $j = 0, 1, \ldots$ **do**

$\beta = 2^j\beta_{t-1}$, $\boldsymbol{z} = \Pi_{\mathbb{S}}(\boldsymbol{v}^{(t)} - \frac{1}{\beta}\nabla g(\boldsymbol{v}^{(t)}))$

**if** $g(\boldsymbol{z}) \leq \tilde{g}_{\beta,\boldsymbol{v}^{(t)}}(\boldsymbol{z})$ **then**

update $\beta_t = \beta$ and $\boldsymbol{z}^{(t+1)} = \boldsymbol{z}$

break

**end if**

**end for**

update $\delta_t = \frac{1+\sqrt{1+4\delta_{t-1}^2}}{2}$

**until** $\boldsymbol{z}^{(t)}$ **converges**;

$\boldsymbol{z}_i = \boldsymbol{z}^{(t)}$.

**end for**

**Output:** LAE vectors $\{\boldsymbol{z}_i\}_{i=1}^n$

#### 1.3 OCAT integration of multiple scRNA-seq datasets

To integrate $H$ single-cell RNA-seq datasets with OCAT, suppose each dataset is of size $N_h$ for $h \in \{1, \ldots, H\}$, and all datasets share $M$ common genes. After the pre-processing and dimension reduction steps, OCAT obtains $X_h \in \mathbb{R}^{N_h \times d}$ as the reduced gene expression matrix for the $h$th dataset.

OCAT then identifies $m_h$ ghost cells from each $X_h$ with features $\{\boldsymbol{u}_k\}_{k=1}^{m_h}$. For the $i$th individual cell, OCAT identifies $s_h$ closest ghost cells with indices $\langle ih \rangle$ from the $h$th ghost cell set for $h \in \{1, \ldots, H\}$. Within the $h$th ghost cell set, OCAT obtains the optimized weights $\boldsymbol{z}_{\langle ih \rangle}$ such that

$$\sum_{k=1}^{m_h} Z_{ik} = 1, \text{ and } Z_{ik} \geq 0 \text{ for } k = 1, 2, \ldots, m_h.$$

Similarly, we obtain the optimized weights for all the ghost cell sets for $h \in \{1, \ldots, H\}$. The weights of the edges connecting the $i$th single cell to all the ghost cells are then denoted as $Z_{i,\langle i \rangle} = (\boldsymbol{z}_{\langle i1 \rangle}, \boldsymbol{z}_{\langle i2 \rangle}, \ldots, \boldsymbol{z}_{\langle iH \rangle})^T$ and $Z_{i,\overline{\langle i \rangle}} = 0$. The collection of all the edge weights of $N$ single cells connecting to $M$ ghost cells is denoted as $Z = \{\boldsymbol{z}_i\}_{i=1}^N \in \mathbb{R}^{N\times M}$, where $N = \sum_{h=1}^H N_h$ and $M = \sum_{h=1}^H m_h$.

Following the message passing procedures outlined in Online Methods Section 5.4.3, we obtain the refined embeddings, $Z_W^{\text{comb}}$, for each single cell through message passing between the combined ghost cells. We lastly normalize $Z_W^{\text{comb}}$ by

$$Z_W^{\text{comb, norm}} = \frac{Z_W^{\text{comb}}}{||Z_W^{\text{comb}}||}.$$

### 2 Evaluation metrics

#### 2.1 Similarity metric between single cell and "ghost" cell

We use cosine similarity as the similarity measure to identify the $s$ closest "ghost" cells in constructing the sparse bipartite graph. The cosine similarity between single cell $i$ and "ghost" cell $j$ is defined as

$$\theta_{ij} = \frac{\boldsymbol{y}_i^T \boldsymbol{u}_j}{\sqrt{\boldsymbol{y}_i^T \boldsymbol{y}_i \boldsymbol{u}_j^T \boldsymbol{u}_j}},$$

where $\boldsymbol{y}_i$ is the feature of the $i$th single cell and $\boldsymbol{u}_j$ is the feature of the $j$th "ghost" cell.

#### 2.2 Metrics for evaluating cell type clustering performance

To assess the cell type clustering performance of the OCAT features, we compute the normalized mutual information (NMI) score of the predicted cell types and the annotated cell type labels ("ground truth"). The NMI score measures the overlap between the predicted cell type labels and true cell type labels normalized by entropy. We define entropy as

$$H(X) = -\sum_{i=1}^{|X|} \frac{|X_i|}{N} \log \frac{|X_i|}{N}, \tag{4}$$

where $|X|$ is the total number of categories, $|X_i|$ is the number of observations fall into category $i$ and $N$ is the total number of observations.

The NMI score of cell type clustering is defined as

$$\mathrm{NMI}_{\text{cluster}}(U, V) = \frac{2\mathrm{MI}(U, V)}{H(U) + H(V)}, \quad \text{where } \mathrm{MI}(U, V) = \sum_{i=1}^{|U|} \sum_{j=1}^{|V|} \frac{|U_i \cap V_j|}{N} \log \frac{N|U_i \cap V_j|}{|U_i||V_j|}, \tag{5}$$

$U$ is the set of sets of predicted cell types, $V$ is the set of sets of annotated cell types ("ground truth"), $N$ is the total number of cells, and $H(\cdot)$ is the entropy defined in (4). The value of $\mathrm{NMI}_{\text{cluster}}$ ranges from 0 to 1, where $\mathrm{NMI}_{\text{cluster}} = 1$ indicates perfect clustering of single cells by the annotated cell types, and $\mathrm{NMI}_{\text{cluster}} = 0$ indicates random guessing. In this work, we adopt the `normalized_mutual_info_score` function in the `scikit-learn` Python package [2] to compute the NMI scores.

#### 2.3 Metrics for evaluating batch effect correction

OCAT does not impose explicit batch correction steps, but we demonstrate that OCAT is robust to batch effect by capturing the global similarities between single cells. To assess the batch effect correction performance of OCAT, we adopt a modified NMI metric as in Section 2.2, and define $\mathrm{NMI}_{\text{batch}}$ as

$$\mathrm{NMI}_{\text{batch}}(I, J) = \frac{2\mathrm{MI}(I, J)}{H(I) + H(J)}, \tag{6}$$

where $I$ is the predicted batch labels, $J$ is the true batch origin of each single cell, and MI is defined as in (5). $\mathrm{NMI}_{\text{batch}} = 0$ indicates that the predicted labels are not confounded with the batch label, and $\mathrm{NMI}_{\text{batch}} = 1$ indicates that the predicted labels are perfectly confounded with the batch labels. We report $(1 - \mathrm{NMI}_{\text{batch}})$ such that a higher $(1 - \mathrm{NMI}_{\text{batch}})$ value indicates better batch effect removal performance.

#### 2.4 Other variations of clustering performance evaluation metrics

Besides the NMI score, we also assess the cell type clustering and batch effect correction performance using the adjusted mutual information (AMI) scores. The AMI score is defined as

$$\mathrm{AMI}(U, V) = \frac{\mathrm{MI}(U, V) - E(\mathrm{MI}(U, V))}{\mathrm{mean}(H(U), H(V)) - E(\mathrm{MI}(U, V))}, \tag{7}$$

where $U$ is the predicted label; $V$ is the ground truth label; $E(\cdot)$ is the expectation; $H$ is the entropy defined in (4); MI is defined as in (5). We compute the AMI scores using the `adjusted_mutual_info_score` function in the `scikit-learn` Python package [2].

The rand index (RI) is a measure of the percentage of correct decisions based on the true positives ($tp$), false positives ($fp$), true negatives ($tn$), and false negatives ($fn$) counts. The RI is defined as

$$\mathrm{RI} = \frac{tp + tn}{tp + fp + fn + tn}. \tag{8}$$

Here we also adopt the adjusted rand index (ARI) to assess the cell type clustering and batch effect correction. The ARI is defined as,

$$\mathrm{ARI} = \frac{\mathrm{RI} - E(\mathrm{RI})}{\max(\mathrm{RI}) - E(\mathrm{RI})}, \tag{9}$$

where RI is the rand index defined in (8) and $E(\cdot)$ is the expectation. We compute the ARI scores using the `adjusted_rand_score` function in the `scikit-learn` Python package [2].

We report the $\mathrm{AMI}_{\mathrm{cluster}}$, $1 - \mathrm{AMI}_{\mathrm{batch}}$, $\mathrm{ARI}_{\mathrm{cluster}}$, and $1 - \mathrm{ARI}_{\mathrm{batch}}$ metrics for all benchmarking integration datasets in Table S1. We also report the AMI and ARI scores for clustering individual datasets in Table S2.

#### 2.5 Runtime and memory usage

We benchmarked the runtime and memory usage of OCAT against Seurat v3 [23], Harmony [7] and Scanorama [6]. We timed the function call that performs data integration using the `time.time()` function in Python. We monitored the memory usage with HTOP and recorded the maximum memory usage occurred during the execution of each integration pipeline on a desktop workstation (Intel(R) Xeon(R) CPU @ 3.60GHz processor).

#### 2.6 Evaluation metric of trajectory and pseudotime inference

We benchmark OCAT with Slingshot [22], PAGA Tree [29] and Monocle ICA [17] on trajectory and pseudotime inference performance. We adopted the benchmarking procedure in dynverse [19]. Dynverse defines five categories (i.e. features, cell positions, neighbourhood, topology, branch assignment) with 16 metrics to evaluate the performance of trajectory and pseudotime inference; see Table S7 for the details of these metrics.

For the HSMM dataset, we also assessed the accuracy of OCAT pseudotime inference by reporting the Pearson correlation between the OCAT predicted pseudotime with the real time stamp labels by cell type cluster. The Pearson correlation $r$ is defined as:

$$r = \frac{\sum_{i=1}^{k}(u_i - \bar{u})(v_i - \bar{v})}{\sqrt{\sum_{i=1}^{k}(u_i - \bar{u})^2 \sum_{i=1}^{k}(v_i - \bar{v})^2}}, \tag{10}$$

where $u_i$ is the average inferred pseudotime of cells in cluster $i$, and $v_i$ is the average real time stamp of cells in cluster $i$, with $i \in \{1, 2, \ldots, k\}$ clusters. $\bar{u}$ is the average predicted pseudotime, and $\bar{v}$ is the average real time stamp. We used the `pearsonr` function in the `scipy.stats` Python package [2] to calculate the Pearson correlation score $r$.

##### 2.7 Cell inference evaluation metrics

To assess the performance of cell type assignment in cell inference, we used the *precision*, *recall* and $f1$ scores implemented in Python in the `scikit-learn` package [2]. We calculated the average of *precision*, *recall* and $f1$ across all classes, weighted by number of samples in each class; see details in Table S5. The *precision*, *recall* and $f1$ scores for each class are calculated from true positives ($tp$), false positives ($fp$), true negatives ($tn$), and false negatives ($fn$) in that particular class.

$$precision = \frac{tp}{tp + fp}, \quad recall = \frac{tp}{tp + fn} \text{ and } f1 = \frac{2 \times precision \times recall}{precision + recall}. \tag{11}$$

### 3 Datasets

#### 3.1 Integration datasets

We evaluated OCAT's integration performance on five multi-source scRNA-seq datasets from [24]. Each dataset contains two or more batches of scRNA-seq data from different experiments or sequencing technologies for similar cell classes, presenting five different scenarios for batch correction.

- **Human dendritic dataset**: contains two batches of scRNA-seq data sequenced using Smart-Seq2, with four cell populations identified, namely CD1C DC, CD141 DC, plasmacytoid DC, and double negative cells [26]. [24] further removed cell type CD1C DC from batch 1 and cell type CD141 DC from batch 2, to create non-identical cell types among the batches. The resulting batch 1 contains 288 cells with three annotated cell types CD141 DC, double negative, plasmacytoid DC; batch 2 also contains 288 cells with three annotated cell types CD1C DC, double negative, plasmacytoid DC. The two batches share two common cell types plasmacytoid DC and double negatives, with one unshared cell type respectively (CD141 DC and CD1C DC). This dataset presents a common scenario of batch effect with the presence of non-identical cell types.

- **Mouse atlas dataset**: contains two batches of scRNA-seq generated independently by [5] using Microwell-Seq and the Tabula Muris Consortium [3] with 10x Genomics and Smart-Seq2 protocols. The 11 cell types with the highest cell numbers in both batches were retained. The resulting first batch contains the read counts of 4239 cells and the second batch contains 2715 cells, with 15006 common genes. This dataset captures the batch effect resulting from different sequencing technologies.

  The mouse atlas dataset contains the following 11 annotated cell types: T-cell, stromal, endothelial, macrophage, monocyte, epithelial, B-cell, neutrophil, dendritic, smooth-muscle, NK. All cell types present in both batches.

- **Human pancreas dataset**: The human pancreas dataset consists of data from five different sources ([1], [20], [10], [28], [30]) of human pancreatic cells. [24] further processed the dataset by removing cells with ambiguous annotations, and the resulting batches contain a total of 14767 cells with 15 different cell types. This dataset captures the batch effect across multiple sequencing technologies.

  In the resulting five batches, [1] contains 8569 cells of 13 annotated cell types (acinar, beta, delta, stellate, ductal, alpha, epsilon, gamma, endothelial, macrophage, schwann, mast, t cell); [10] contains 2122 cells of 9 annotated cell types (alpha, ductal, endothelial, delta, acinar, beta, gamma, mesenchymal, epsilon); [20] contains 2127 cells of 11 annotated cell types (delta, alpha, gamma, ductal, acinar, beta, MHC, stellate, endothelial, epsilon, mast); [28] contains 457 cells of 7 annotated cell types (alpha, ductal, delta, beta, gamma, acinar, mesenchymal); [30] contains 1492 cells of 4 annotated cell types (beta, alpha, delta, gamma).

- **PBMC dataset**: The PBMC dataset contains two batches of PBMC data from healthy donors generated by 3' and 5' 10x Genomics protocols respectively. 8098 cells from the 3' batch and 7378 cells from the 5' batch were selected and annotated by [16] with k-nearest neighbor clustering based on canonical markers. This dataset presents the challenge of integrating data with biological differences caused by sequencing protocols.

  The PBMC dataset contains 9 annotated cell types: CD4 T cell, CD8 T cell, monocyte CD14, B cell, NK cell, monocyte FCGR3A, plasmacytoid dendritic cell, megakaryocyte, hematopoietic stem cell. All cell types exist in both batches.

- **Mouse hematopoietic dataset**: The mouse hematopoietic dataset contains two batches of mouse hematopoietic stem and progenitor cells, generated by [12] with SMART-seq2 protocol and by [13] with MARS-seq protocol respectively. [24] further extracted 2729 well-annotated cells from the MARS-seq dataset. The resulting SMART-seq2 data contains 1920 cells and MARS-seq data contains 2729 cells, with 3467 genes common genes retained.

  The MARS-seq batch contains 3 annotated cell types: GMP, MEP, and CMP. The SMART-seq2 batch contains 7 annotated cell types: MPP, MEP, CMP, LTHSC, LMPP, Unsorted, and GMP. This dataset therefore captures the batch effect from not only different sequencing technologies but also non-identical cell types.

#### 3.2 Individual datasets

- **The Romanov dataset** [18] contains the RNA-seq data from 2881 single cells with 24341 genes in mouse hypothalamus obtained using Illumina HiSeq platform. Seven cell type clusters were identified by divisive biclustering method and annotated based on lineage-specific protogene markers.

  The Romanov dataset contains 7 annotated cell types: oligos, neurons, ependymal, astrocytes, endothelial, vsm, microglia.

- **The Zeisel dataset** [31] contains the 3'-end counts of the unique molecular identifiers (UMI) assays from 3005 single cells with 4412 genes in the mouse somatosensory cortex and hippocampal CA1 region.

  The Zeisel dataset contains 9 annotated cell types: S1 and CA1 pyramidal neurons, interneurons, oligodendrocytes, astrocytes, microglia, vascular endothelial cells, mural cells, and ependymal cells.

- **The mouse retina dataset** [21] contains 19,829 single cells with 13,166 genes extracted from the Macoskco data set (originally 44,808 cells), containing UMI (3'-end) counts obtained by Drop-seq.

  The Mouse Retina dataset contains 15 annotated cell types: BC1A, BC1B, BC2, BC3A, BC3B, BC4, BC5A, BC5B, BC5C, BC5D, BC6, BC7, BC8_9, MG, RBC.

- **The PBMC 68k dataset** [32] contains the UMI (3'-end) counts from 68579 peripheral blood mononuclear cells (PBMCs) with 1000 genes profiled using 10x Genomics GemCode platform.

  The PBMC 68k dataset contains 11 annotated cell types: CD14+ Monocyte, CD19+ B, CD34+, CD4+ T Helper2, CD4+/CD25 T Reg, CD4+/CD45RA+/CD25- Naive T, CD4+/CD45RO+ Memory, CD56+ NK, CD8+ Cytotoxic T, CD8+/CD45RA+ Naive Cytotoxic, Dendritic.

- **The human skeletal muscle myoblast (HSMM) dataset** [25] contains time-series RNA-seq data of 271 cells collected at 0, 24, 48 and 72 hours since human myoblast culture in the differentiation media. Five cell groups were annotated using GSVA based on known gene markers: myoblasts, intermediates, myotubes, fibroblasts, and undifferentiated.

### 4 Hyperparameter selection and sensitivity analysis

The sparse encoding workflow of OCAT involves the specification of three hyperparameters: $d$ as the projection dimension from the gene feature space by online FSM in the pre-processing step, $m$ as the number of "ghost" cells each single cell connects to in the bipartite graph, and $s$ as the closest "ghost" cells to construct the sparsified encoding of each cell. Here we define $p = s/m$ as the percentage of closet "ghost" cells chosen to embed the gene expression of each single cell. The number of closest "ghost" cells is computed as $s = \lceil pm \rceil$. See the Online methods Section for details on $d$, $m$ and $s$.

Here we performed a set of sensitivity analyses to assess the impact of hyperparameter values on the OCAT sparse encoding in integrating multiple scRNA-seq data as well as clustering individual scRNA-seq data. We demonstrate that the OCAT sparse encoding is robust to various combinations of hyperparameters by comparing the cell type clustering performance.

#### 4.1 Hyperparameter sensitivity analysis in integrating multiple datasets

We used the Mouse Atlas dataset as an example to demonstrate the robustness of OCAT embedding in integrating multiple datasets. We reported the NMI of cell type clustering using OCAT embeddings with $m_1 = m_2 = 45$, $d = 70$ and $p = 0.3$ in Table 1, where $m_1$ and $m_2$ are the number of "ghost" cells selected for batch 1 and batch 2. We first set $m_1 = m_2 = m$, and assessed the sensitivity of $d$, $m$ and $p$ on the cell type clustering performance by taking the following hyperparameter values: $d \in \{40, 50, 60, 70, 80, 90, 100, 110, 120\}$; $m \in \{20, 25, 30, 35, 40, 45, 50, 55, 60\}$; $p \in \{0.1, 0.3, 0.5, 0.7\}$. Figure S4A plots the $\text{NMI}_{\text{cell type}}$ of each hyperparameter combination. The NMI values range from 0.7286 to 0.8007 with median 0.7809 and standard deviation 0.0144, which implies that the OCAT encoding is robust to the specification variations of $d$, $m$ and $p$ in integrating multiple datasets.

We then assessed the impact of values of $m_1$ and $m_2$ on the OCAT sparse encoding. We allow $m_1$ and $m_2$ to be different, taking values $m_1, m_2 \in \{20, 25, 30, 35, 40, 45, 50, 55, 60\}$ while fixing $d = 70$ and $p = 0.3$. Figure S4C plots the $\text{NMI}_{\text{cell type}}$ of each hyperparameter combination with a heatmap. The NMI values range from 0.7378 to 0.8007 with median 0.7812 and standard deviation 0.0162, which implies that the OCAT encoding is robust to the specification of the number of "ghost" cells in each batch.

#### 4.2 Hyperparameter sensitivity analysis in single datasets

We next assessed the sensitivity of $d$, $m$ and $p$ on clustering performance using the Zeisel dataset as an example. We reported the NMI of cell type clustering using OCAT embeddings with $m = 50$, $d = 30$ and $p = 0.3$ in Table 2. Here, we assessed the sensitivity by taking the following hyperparameter values $d \in \{20, 30, 40, 50, 60, 70, 80, 90, 100\}$, $m \in \{30, 35, 40, 45, 50, 55, 60, 65, 70\}$ and $p \in \{0.1, 0.3, 0.5, 0.7\}$. Figure S4B plots the cell type clustering NMI metrics of each hyperparameter combination. Under each combination, the variation of the NMI metrics is minimal, where the NMI values range from 0.6572 to 0.7987 with median 0.7591 and standard deviation 0.0152. This demonstrates that the OCAT sparse encoding is robust to hyperparameter specifications in single datasets.

#### 4.3 Hyperparameter recommendations

The OCAT package allows the users to specify their desired values for $d$, $m$ and $p$ as input arguments. We recommend the user select $d$ based on the number of genes in the scRNA-seq expression matrices, with larger dimension $d$ corresponding to larger number of genes. We recommend the choice of $m$, the number of “ghost” cells, based on the number of single cells, with larger $m$ to encode larger number of single cells. We recommend the choice of $p$ based on the sparsity of the desired encoding. By default, $p$ is set to 0.3.

### 5 Differential gene analysis and comparison

OCAT effectively identifies the differential genes for cell type annotation. We demonstrate the performance of OCAT differential gene analysis using the Zeisel dataset [31]. We summarize the top 5 OCAT selected differential genes in Table S6, and show that OCAT manages to replicate the marker genes reported by [31]. For example, *Gad1* and *Gad2* genes for Interneuron cells, and *Acta2* gene for mural cells.

We further compare the top differential genes for each cell type identified by OCAT and Seurat, and report the number of top differential genes selected by both methods. Table S6 summarizes the top 5 differential genes identified by both OCAT and Seurat, and the identified genes are highly consistent for most cell types. We also compare the top $k$ differential genes identified by the two methods, and plot the number of overlapped genes ($\leq k$) against $k$, the number of top differential genes, in Figure S6. The closer a dot to the diagonal line, the more overlap between the top differential genes selected by both methods. The plotted line for each cell type is close to the dotted diagonal line, which suggests that the top differential genes selected by OCAT and Seurat are highly consistent.

### 6 Benchmarking specifics

#### 6.1 Integrating multiple datasets

We benchmark the cell type clustering performance and batch correction performance of OCAT against Seurat v3 [23], Scanorama [6] and Harmony [7]. We followed the recommended settings and workflows of all the methods, with the specific parameters as follows. For Seurat v3, we followed the default settings, with no cells or genes screened out and selected 2,000 most variable genes. We chose the top 50 principal components as the Seurat embeddings for the cells. To benchmark against Harmony, we adopted the Seurat object pre-processed as in the Seurat v3 benchmarking process, but extracted the top 50 features as the Harmony features using the `RunHarmony` function. For Scanorama, we added a manual step to normalize the raw scRNA-seq count matrix, and ran its recommended pipeline. We subsequently performed k-means clustering on the embeddings using the `KMeans` function, and computed the NMI, AMI and ARI for batch correction and cell type clustering metrics using the `adjusted_rand_score`, `adjusted_mutual_info_score`, `normalized_mutual_info_score` functions from the `sklearn` python package.

#### 6.2 Individual scRNA-seq datasets

We benchmark the cell type clustering performance of OCAT with Seurat v3 [23], scVI [9] and SIMLR [27]. For Seurat v3, we followed the default settings, with no cells screened out in the gene by cell count matrix. We chose the top 50 principal components as the Seurat features of the single cells. We adopt the Seurat default `FindClusters` function to cluster the cells. When benchmarking with scVI (version 0.6.5), we followed the default settings, and down-sampled 500 most variable genes out of all the genes available. We set the maximum number of epochs as 400, and used 90% of the dataset as the training set. We lastly used k-means to predict the clusters of the cells. We also benchmarked OCAT with SIMLR. SIMLR takes the same raw cell by gene matrix as OCAT. We adopted the `SIMLR_Large_Scale` function for all datasets. For the Zeisel dataset, we transformed the gene expression matrix $X$ to $\log(X+1)$, and set the SIMLR tuning parameter $k = 50$ and assessed number of principal components $kk = 150$. For the PBMC dataset, we used the raw cell by gene matrix and adopted the default SIMLR settings $k = 10$ and $kk = 100$. For the retina dataset, we used the raw cell by gene matrix and set the tuning parameter $k = 30$ and $kk = 300$. For the Romanov dataset, we used the raw cell by gene matrix and set the tuning parameter $k = 25$ and $kk = 400$. We performed k-means clustering on the embeddings using the `KMeans` function, and computed the NMI, AMI and ARI for cell type clustering metrics using the `adjusted_rand_score`, `adjusted_mutual_info_score`, `normalized_mutual_info_score` functions from the `sklearn` python package.

#### 6.3 Benchmarking OCAT trajectory and pseudotime inference

We benchmarked OCAT's trajectory and pseudotime inference performance with Slingshot [22], PAGA Tree [29] and Monocle ICA [17] using the `dynverse` R package on 28 gold-standard real datasets [19]. The input datasets consist of expression matrices from a wide range of dynamic processes and sequencing technologies. The gold-standard annotations on trajectory topologies were obtained from biological evidence besides the expression matrices, including four different trajectory types (i.e. linear, tree, multifurcation, bifurcation). We evaluated the trajectory and pseudotime assignment by each method using five aggregated metrics (features, cell positions, neighbourhood, topology, branch assignment) derived from 16 dynverse metrics; see Supplementary Table S7 for details on metric aggregation and interpretation.

In OCAT's benchmarking workflow, for each dataset, we first extracted the OCAT sparse encodings of the single cells, with hyperparameters $d$ by package default and $m$ selected based on the number of single cells.

We subsequently inferred cluster labels by k-means clustering on the sparse encodings and then ran OCAT trajectory analysis to estimate the cell lineages. For pseudotime analysis, we used the root cluster inferred by OCAT in the previous trajectory step as input. For Slingshot [22], PAGA Tree [29] and Monocle ICA [17], we used the benchmarking pipeline provided by [19] with dynverse specified hyperparameters, which computes the lineages and pseudotime of the single cells in the reduced feature space for each dataset in a similar fashion.

### 7 Data and code availability

All the scRNA-seq data analyzed are publicly available, and were retrieved from `https://hub.docker.com/repository/docker/jinmiaochenlab/batch-effect-removal-benchmarking` and `https://www.baderlab.org/Software/Tempora`.

OCAT is freely available at `https://github.com/bowang-lab/OCAT`.

| Dataset | metric | OCAT | Seurat v3 | Harmony | Scanorama |
|---|---|---|---|---|---|
| Human dendritic [26] | $NMI_{cell\ type}$ | **0.7718** | 0.7375 | 0.7653 | 0.7212 |
| | $1\text{-}NMI_{batch}$ | 0.9999 | 0.9999 | **1.0000** | 0.9999 |
| | $AMI_{cell\ type}$ | **0.7703** | 0.7357 | 0.7637 | 0.7194 |
| | $1\text{-}AMI_{batch}$ | 1.0012 | 1.0012 | **1.0013** | 1.0012 |
| | $ARI_{cell\ type}$ | **0.7533** | 0.6932 | 0.7350 | 0.7188 |
| | $1\text{-}ARI_{batch}$ | 1.0014 | 1.0014 | **1.0015** | 1.0014 |
| Mouse atlas | $NMI_{cell\ type}$ | **0.8006** | 0.6981 | 0.7625 | 0.6960 |
| | $1\text{-}NMI_{batch}$ | **0.9999** | 0.9970 | 0.9959 | 0.9905 |
| | $AMI_{cell\ type}$ | **0.8000** | 0.6971 | 0.7617 | 0.6950 |
| | $1\text{-}AMI_{batch}$ | **1.0000** | 0.9971 | 0.9960 | 0.9906 |
| | $ARI_{cell\ type}$ | **0.7506** | 0.5057 | 0.6487 | 0.5778 |
| | $1\text{-}ARI_{batch}$ | 0.9991 | 1.0108 | **1.0122** | 0.9751 |
| Human pancreas | $NMI_{cell\ type}$ | **0.7949** | 0.7947 | 0.7302 | 0.7249 |
| | $1\text{-}NMI_{batch}$ | 0.9650 | **0.9762** | 0.9600 | 0.9538 |
| | $AMI_{cell\ type}$ | **0.7943** | 0.7940 | 0.7293 | 0.7241 |
| | $1\text{-}AMI_{batch}$ | 0.9654 | **0.9767** | 0.9605 | 0.9542 |
| | $ARI_{cell\ type}$ | 0.6455 | **0.7278** | 0.5876 | 0.4332 |
| | $1\text{-}ARI_{batch}$ | 1.0131 | 1.0435 | **1.0585** | 1.0109 |
| PBMC | $NMI_{cell\ type}$ | 0.7424 | **0.7932** | 0.7851 | 0.7141 |
| | $1\text{-}NMI_{batch}$ | 0.9934 | 0.9943 | **0.9944** | 0.9939 |
| | $AMI_{cell\ type}$ | 0.7421 | **0.7929** | 0.7848 | 0.7137 |
| | $1\text{-}AMI_{batch}$ | 0.9934 | 0.9943 | **0.9945** | 0.9940 |
| | $ARI_{cell\ type}$ | 0.6239 | **0.7064** | 0.7005 | 0.5627 |
| | $1\text{-}ARI_{batch}$ | 0.9905 | 0.9908 | **0.9910** | 0.9904 |
| Mouse hematopoietic | $NMI_{cell\ type}$ | 0.5019 | 0.4606 | 0.4111 | **0.5160** |
| | $1\text{-}NMI_{batch}$ | 0.9648 | 0.9673 | **0.9734** | 0.9674 |
| | $AMI_{cell\ type}$ | 0.5007 | 0.4594 | 0.4098 | **0.5149** |
| | $1\text{-}AMI_{batch}$ | 0.9649 | 0.9675 | **0.9736** | 0.9676 |
| | $ARI_{cell\ type}$ | 0.5031 | 0.4297 | 0.3727 | **0.5111** |
| | $1\text{-}ARI_{batch}$ | 0.9880 | 0.9977 | **1.0010** | 0.9928 |

Table S1: **OCAT cell type clustering and batch correction performance for integrating multiple scRNA-seq datasets**, compared with Seurat v3, Harmony and Scanorama; see Section 2.2 for details on the evaluation metrics.

| Dataset | metric | OCAT | scVI | Seurat v3 | SIMLR |
|---|---|---|---|---|---|
| Romanov [18] | NMI | **0.6443** | 0.5436 | 0.6343 | 0.4234 |
| | AMI | **0.6429** | 0.5057 | 0.5068 | 0.3993 |
| | ARI | **0.6536** | 0.5010 | 0.4045 | 0.3797 |
| Zeisel [31] | NMI | **0.7884** | 0.7126 | 0.6724 | 0.7373 |
| | AMI | **0.7872** | 0.6505 | 0.5653 | 0.6798 |
| | ARI | **0.7312** | 0.5758 | 0.5369 | 0.5754 |
| Retina [21] | NMI | **0.8742** | 0.7572 | 0.7865 | 0.6337 |
| | AMI | **0.8739** | 0.6865 | 0.6956 | 0.6083 |
| | ARI | **0.6415** | 0.4615 | 0.4332 | 0.5554 |
| PBMC 68k [32] | NMI | **0.5750** | 0.4638 | 0.4899 | 0.5344 |
| | AMI | **0.5749** | 0.4198 | 0.4045 | 0.4304 |
| | ARI | **0.4336** | 0.2764 | 0.2382 | 0.2930 |

Table S2: **OCAT cell type clustering performance for integrating individual scRNA-seq datasets**, compared with scVI, Seurat v3 and SIMLR; see Section 2.2 for details on the evaluation metrics.

| Dataset | $n_{\text{datasets}}$ | $n_{\text{cells}}$ | $n_{\text{genes}}$ | $m$ | $d$ |
|---|---|---|---|---|---|
| Human dendritic [26] | 2 | 576 | 16,594 | 20 | 80 |
| Mouse atlas | 2 | 6, 954 | 5,558 | 45 | 70 |
| Human pancreas | 5 | 14,767 | 15,558 | 65 | 60 |
| PBMC | 2 | 15,476 | 17,430 | 40 | 120 |
| Mouse hematopoietic | 2 | 4,649 | 3,467 | 30 | 70 |

Table S3: **OCAT hyper-parameters for integrating multiple datasets**. $m$ is the number of ghost cells; $d$ is the dimension that OCAT projects the original gene expression matrix to; see Section 3.2 for the detailed description of the datasets.

| Dataset | $n_{\text{cells}}$ | $n_{\text{genes}}$ | $m$ | $d$ |
|---|---|---|---|---|
| Romanov [18] | $2,881$ | $24,341$ | 20 | 30 |
| Zeisel [31] | $3,005$ | $19,972$ | 50 | 30 |
| Retina [21] | $19,829$ | $13,166$ | 60 | 60 |
| PBMC 68k [32] | $68,579$ | $1,000$ | 40 | 40 |

Table S4: **OCAT hyper-parameter settings for individual scRNA-seq datasets**. $m$ is the number of ghost cells; $d$ is the dimension that OCAT projects the original gene expression matrix to; see Section 3.2 for the detailed description of the datasets.

| Dataset | $n_{\text{reference}}$ | $n_{\text{inference}}$ | # of cell types | Precision (weighted) | Recall (weighted) | F1 (weighted) |
|---|---|---|---|---|---|---|
| Romanov [18] | 2,592 | 289 | 7 | 0.8988 | 0.8892 | 0.8854 |
| Zeisel [31] | 2,703 | 302 | 9 | 0.9847 | 0.9834 | 0.9819 |
| Retina [21] | 17,847 | 1,982 | 15 | 0.9879 | 0.9878 | 0.9877 |
| PBMC 68k [32] | 61,721 | 6,858 | 11 | 0.7366 | 0.7696 | 0.7425 |

Table S5: **OCAT cell inference performance** on four individual scRNA-seq datasets: Rovmanov, Zeisel, Retina, and PBMC 68k. In each dataset, $n_{\text{reference}}$ denotes the number of reference cells, and $n_{\text{inference}}$ denotes the number of inference cells. The weighted Precision, Recall and F1 scores between the ground-truth labels and the predicted labels from the inference set are reported to assess the predictive performance.

| Cell Type | OCAT$_{norm}$ | Seurat |
|---|---|---|
| Astrocytes | **1700001C02Rik** | **1700001C02Rik** |
| | Dynlrb2 | 1700009P17Rik |
| | Tmem212 | 1600029I14Rik |
| | **Fam216b** | Gm10714 |
| | Dnali1 | **Fam216b** |
| CA1 | **Crym** | **Crym** |
| | **Cpne6** | Hpca |
| | **Neurod6** | **Gria1** |
| | **Gria1** | **Neurod6** |
| | Wipf3 | **Cpne6** |
| Endothelial | **C1qb** | **C1qa** |
| | **C1qa** | **C1qb** |
| | Fcgr3 | **Tyrobp** |
| | **Fcrls** | **Fcrls** |
| | **Tyrobp** | Fcer1g |
| Ependymal | **Aqp4** | **Acsbg1** |
| | **Acsbg1** | **Lcat** |
| | **Lcat** | **Aqp4** |
| | Gja1 | **Cldn10** |
| | **Cldn10** | Mlc1 |
| Interneurons | **Gad1** | **Gad1** |
| | **Gad2** | **Gad2** |
| | **Slc32a1** | Pnoc |
| | Dlx1 | **Dlx6os1** |
| | **Dlx6os1** | **Slc32a1** |
| Microglia | **Ly6c1** | **Cldn5** |
| | **Cldn5** | Eltd1 |
| | **Flt1** | **Flt1** |
| | Itm2a | **Ly6c1** |
| | Slco1a4 | Pecam1 |
| Mural | **Mog** | **Plp1** |
| | Ugt8a | Trf |
| | Mobp | Mog |
| | Cnp | Mal |
| | Ermn | Apod |
| Oligodendrocytes | **Acta2** | **Acta2** |
| | **Tagln** | Myh11 |
| | **Tpm2** | **Tagln** |
| | **Mustn1** | **Tpm2** |
| | Myl9 | **Mustn1** |
| S1 | **Gm11549** | **Gm11549** |
| | **Mef2c** | **Car10** |
| | **Car10** | Pcsk2 |
| | **Tbr1** | **Tbr1** |
| | Pcp4 | **Mef2c** |

Table S6: **Top five selected differential genes** by OCAT and Seurat v3 in each cell population of the Zeisel dataset.

| Metric | Dynverse Metric | Range (low-high) |
|---|---|---|
| **Features** | Feature importance correlation | $0 - 1$ |
| | Feature importance weighted correlation | $0 - 1$ |
| | Feature importance enrichment ks | $0 - 1$ |
| | Feature importance enrichment wilcox | $0 - 1$ |
| **Cell positions** | Geodesic distance correlation | $0 - 1$ |
| **Neighbourhood** | Random Forest MSE | $0.3 - 0$* |
| | Random Forest normalised MSE | $0 - 1$ |
| | Random Forest $R^2$ | $0 - 1$ |
| | Linear regression MSE | $0.3 - 0$* |
| | Linear regression normalised MSE | $0 - 1$ |
| | Linear regression $R^2$ | $0 - 1$ |
| **Topology** | Edge flip | $0 - 1$ |
| | Hamming-Ipsen-Mikhailov similarity | $0 - 1$ |
| | Isomorphic | $0 - 1$ |
| **Branch assignment** | F1 overlap between the branches | $0 - 1$ |
| | F1 overlap between the milestones | $0 - 1$ |

Table S7: **Evaluation metrics in trajectory and pseudotime inference benchmark.** The aggregated metrics for Features, Cell positions, Neighbourhood, Topology, and Branch assignment are the arithmetic mean of the dynverse metrics in their sub-categories. *The ranges of Random Forest MSE and Linear regression MSE are $0.3 - 0$, and were re-scaled to $0 - 1$ in the calculation of the Neighbourhood metric.

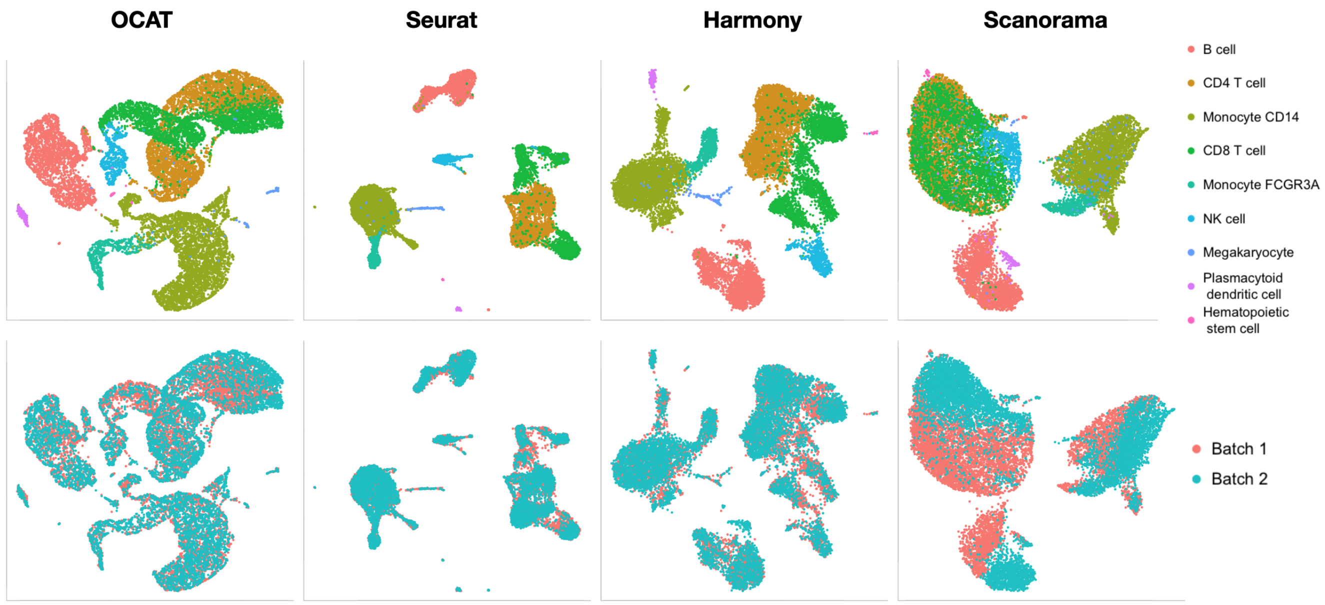

Figure S1: **UMAP projection of the integrated PBMC datasets using OCAT**, benchmarked with Seurat v3, Harmony and Scanorama. The top panels are colored by annotated cell types, and the bottom panels are colored by batch origins.

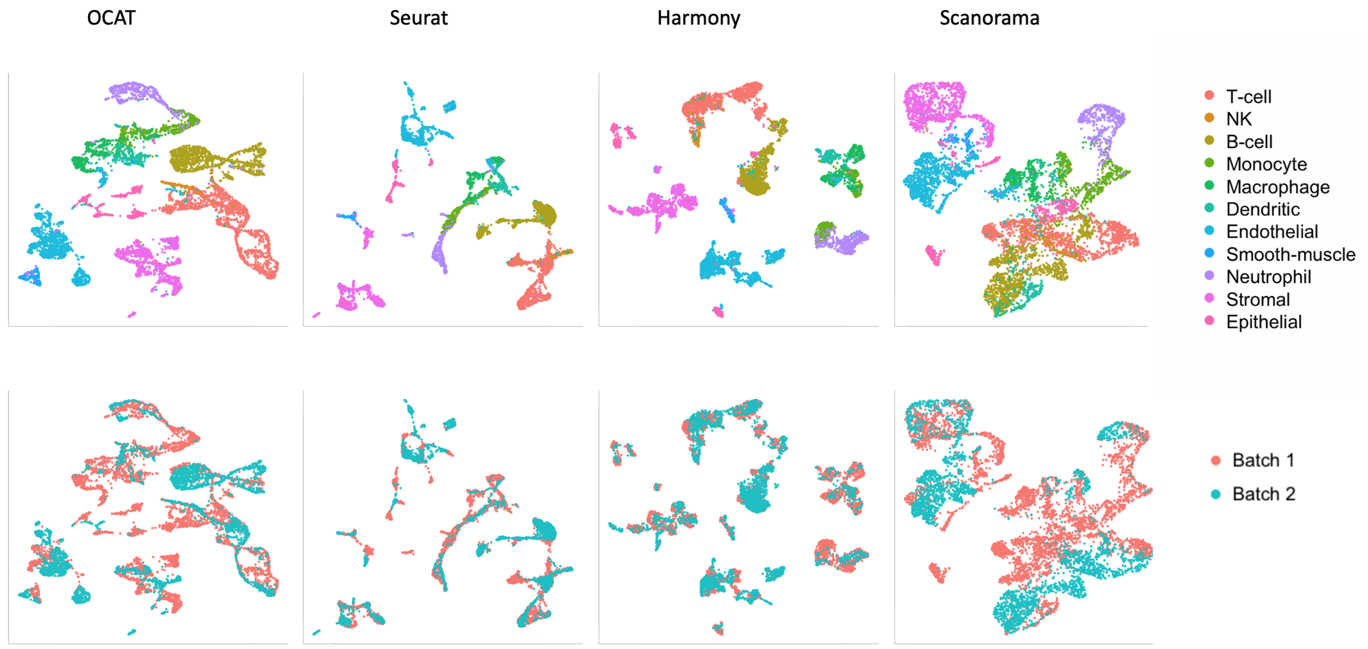

Figure S2: **UMAP projection of the integrated mouse atlas datasets using OCAT**, benchmarked with Seurat v3, Harmony and Scanorama. The top panels are colored by annotated cell types, and the bottom panels are colored by batch origins.

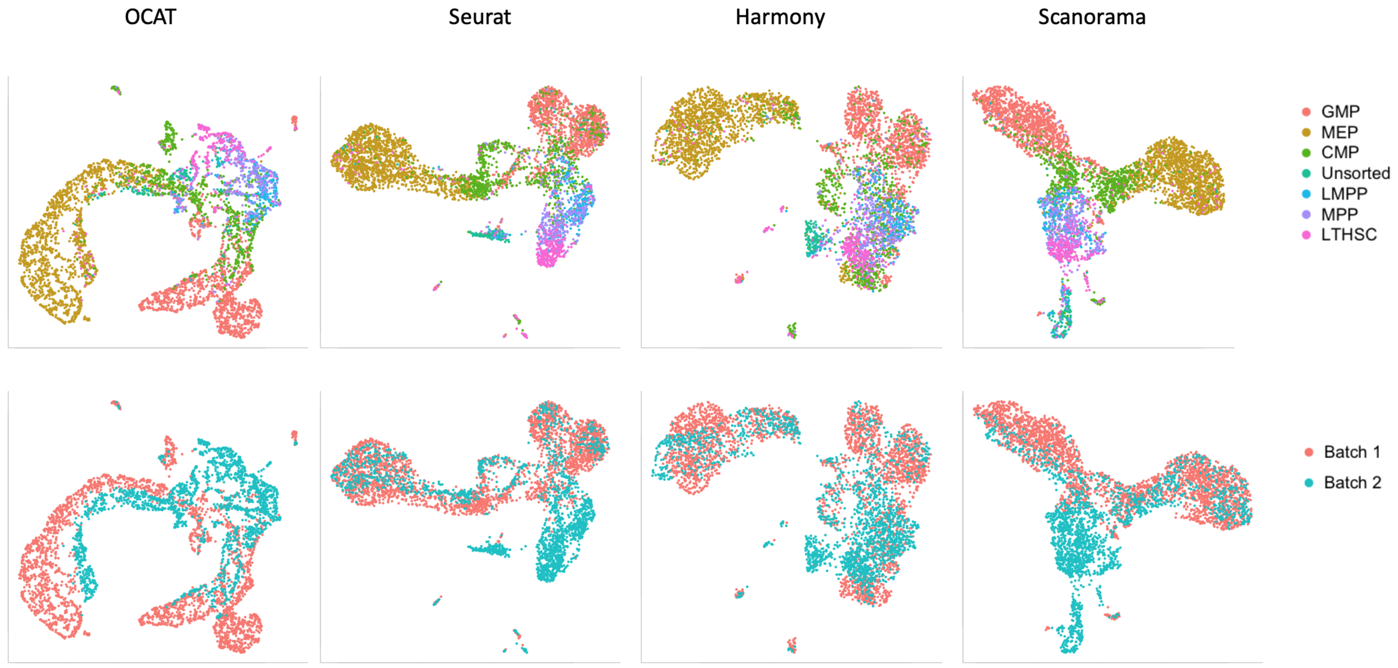

Figure S3: **UMAP projection of the integrated mouse hematopoietic datasets using OCAT**, benchmarked with Seurat v3, Harmony and Scanorama. The top panels are colored by annotated cell types, and the bottom panels are colored by batch origins.

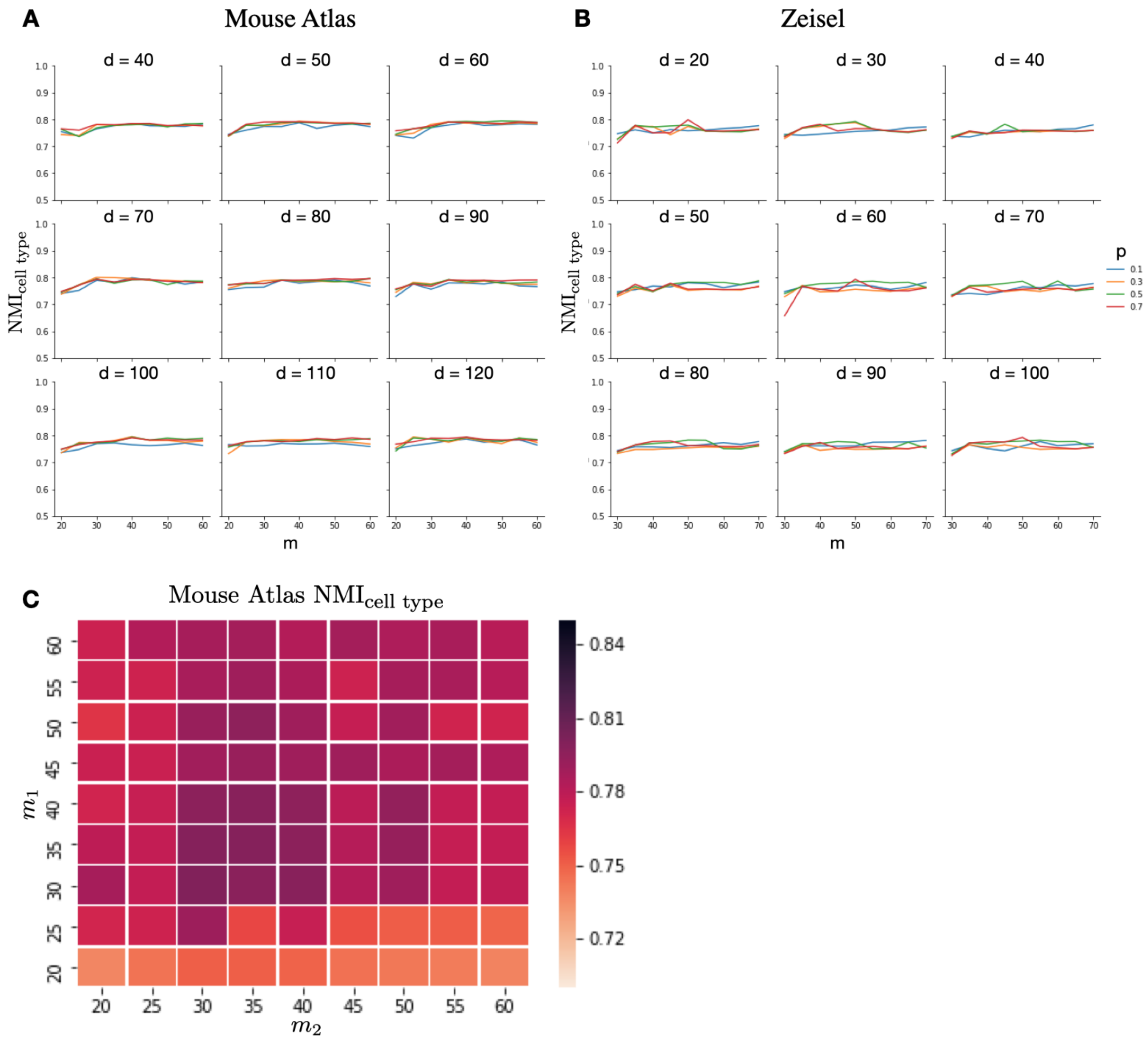

Figure S4: **OCAT hyperparameter sensitivity analysis.** **A**: Cell type clustering performance on integrating two mouse atlas datasets with different hyperparameter values of $d$, $m$, and $p$. The number of ghost cells $m$ in both datasets are set identical. The panel reports $\mathrm{NMI}_{\text{cell type}}$ as the evaluation metric. **B**: Cell type clustering performance on the Zeisel dataset with different hyperparameter values of $d$, $m$, and $p$. The panel reports $\mathrm{NMI}_{\text{cell type}}$ as the evaluation metric. **C**: Heatmap of $\mathrm{NMI}_{\text{cell type}}$ on integrating two mouse atlas datasets with different choices for the number of "ghost" cells (i.e. $m_1$, $m_2$) for the two datasets. $d$ is set to 70 and $p$ is set to 0.3.

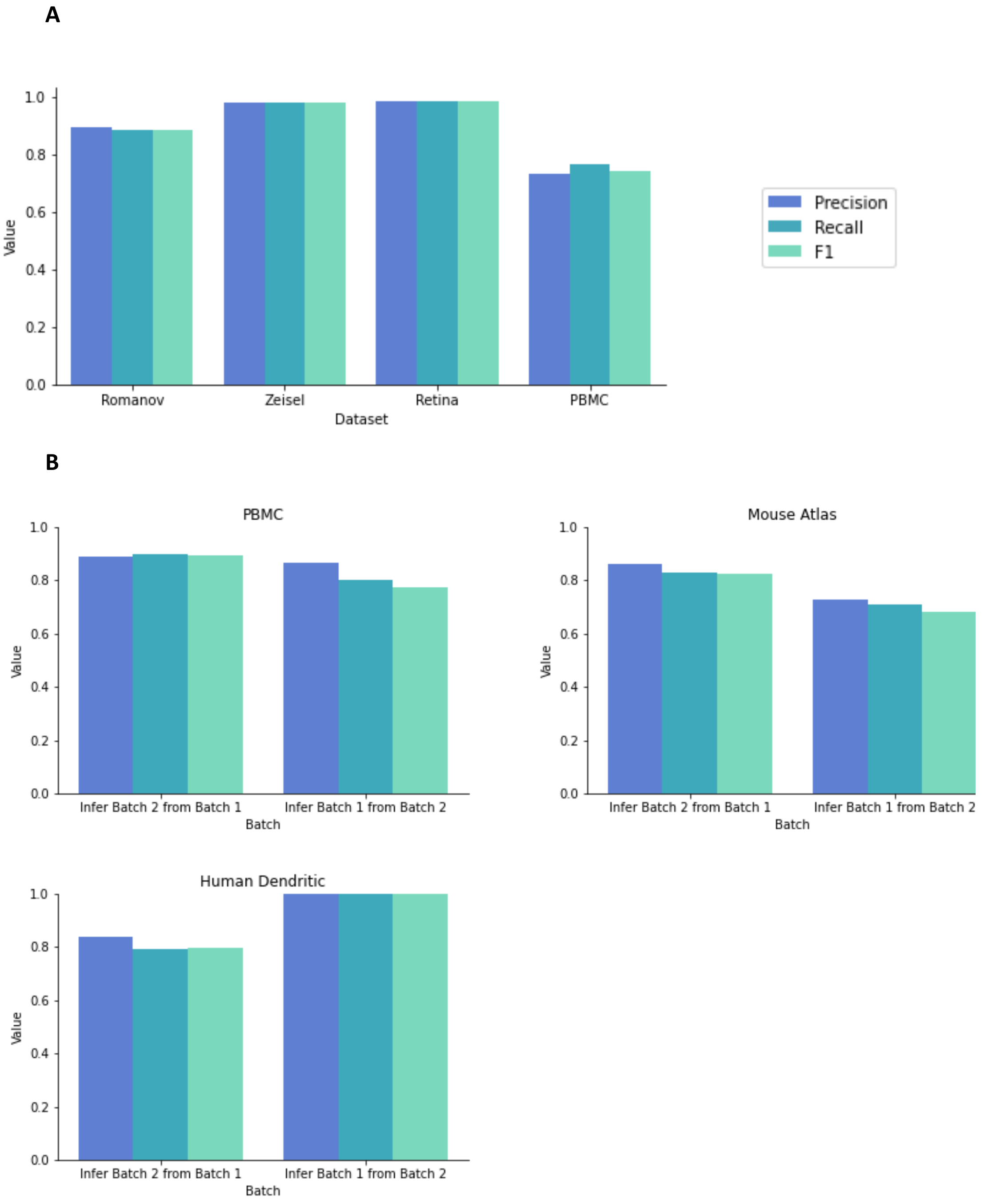

Figure S5: **OCAT cell inference performance on individual and integration datasets.** **A**: OCAT cell type inference performance on individual datasets. The panel reports three classification metrics (i.e. Precision, Recall, F1) on Romanov, Zeisel, Retina and PBMC datasets. **B**: OCAT cell type inference performance on integration datasets. For each of the PBMC, mouse atlas and human dendritic datasets, we reported the Precision, Recall, and F1 scores on two sets of experiments (i.e. infer Batch 2 from Batch 1, infer Batch 1 from Batch 2).

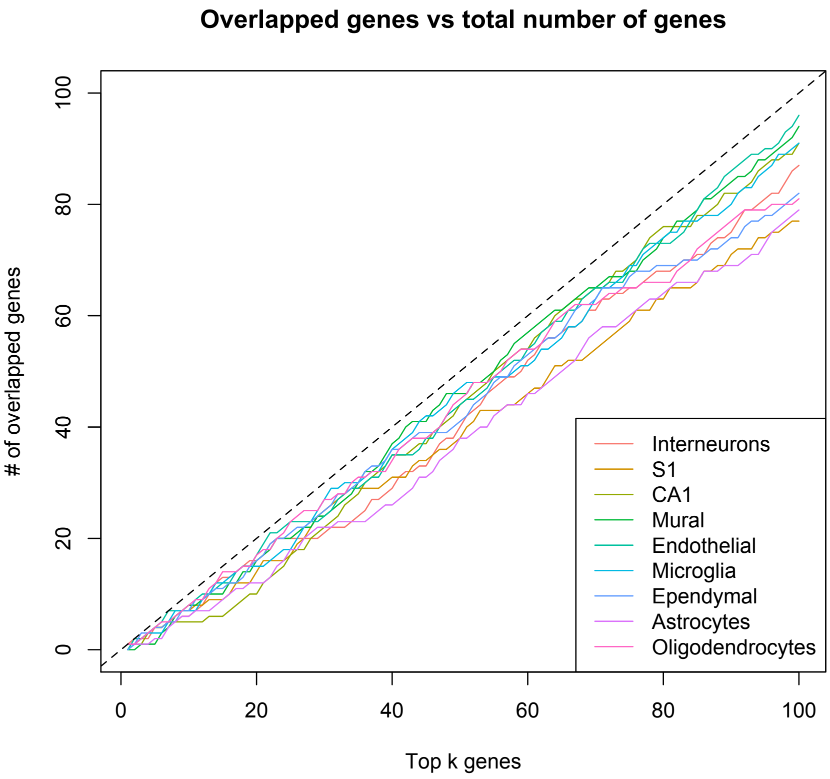

Figure S6: **Top differential gene overlap between OCAT and Seurat in the Zeisel dataset.** The number of top genes, $k$, ranges from 1 to 100. Each colored line represents one cell type group. The dotted diagonal line symbolizes perfect consistency between OCAT and Seurat. The closer a dot is to the diagonal line, the more overlaps between the top $k$ differential genes selected by OCAT and Seurat v3. The top differential genes identified by OCAT and Seurat are mostly consistent for all cell types.

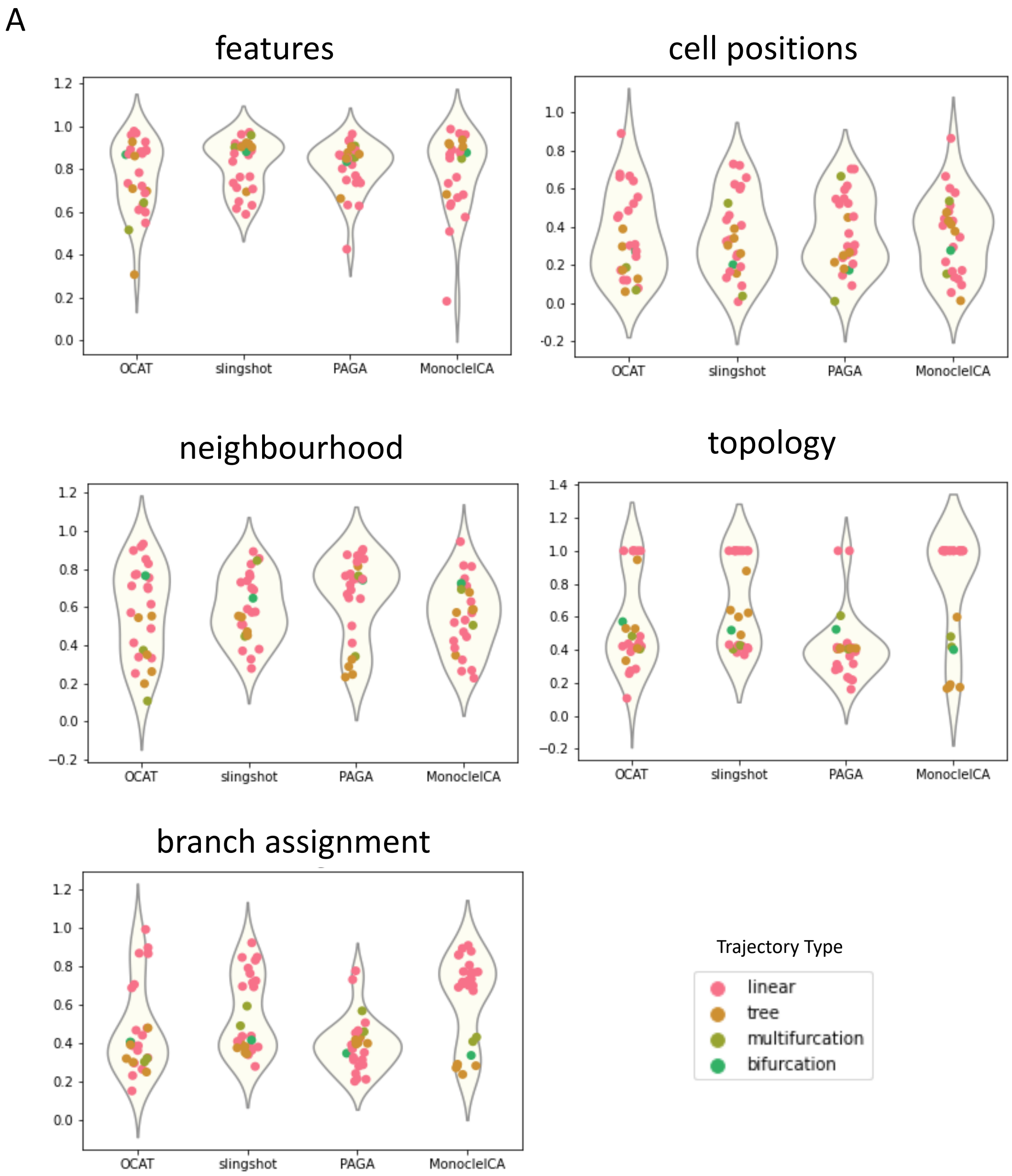

Figure S7: **OCAT trajectory and pseudotime inference benchmark on dynverse real datasets. A**: Violin plot of the OCAT trajectory and pseudotime inference metrics, benchmarking with Slingshot, PAGA and Monocole ICA. Each panel reports the distribution of each of the five aggregated metrics (features, cell positions, neighbourhood, topology, branch assignment) per dataset, color-coded by trajectory type (linear, tree, bifurcation, multifurcation).
